# Serotonin signals through postsynaptic Gα_q_, Trio RhoGEF, and diacylglycerol to promote *C. elegans* egg-laying circuit activity and behavior

**DOI:** 10.1101/2021.05.08.443256

**Authors:** Pravat Dhakal, Sana I. Chaudhry, Rossana Signorelli, Kevin M. Collins

**Affiliations:** Department of Biology, University of Miami, Coral Gables, FL, United States of America

## Abstract

Activated Gα_q_ signals through Phospholipase-Cβ (PLCβ) and Trio, a Rho GTPase exchange factor (RhoGEF), but how these distinct effector pathways promote cellular responses to neurotransmitters like serotonin remains poorly understood. We used the egg-laying behavior circuit of *C. elegans* to determine whether PLCβ and Trio mediate serotonin and Gα_q_ signaling through independent or related biochemical pathways. Our genetic rescue experiments suggest that PLCβ functions in neurons while Trio RhoGEF functions in both neurons and the postsynaptic vulval muscles. While Gα_q_, PLCβ, and Trio RhoGEF mutants all fail to lay eggs in response to serotonin, optogenetic stimulation of the serotonin-releasing HSN neurons restores egg laying only in PLCβ mutants. PLCβ mutants showed vulval muscle Ca^2+^ transients while strong Gα_q_ and Trio RhoGEF mutants had little or no vulval muscle Ca^2+^ activity. Treatment with phorbol 12-myristate 13-acetate (PMA) that mimics 1,2-diacylglycerol (DAG), a product of PIP_2_ hydrolysis, rescued egg-laying circuit activity and behavior defects of Gα_q_ signaling mutants, suggesting both Phospholipase-C and Rho signaling promote synaptic transmission and egg laying via modulation of DAG levels. DAG activates effectors including UNC-13, however we find that phorbol esters, but not serotonin, stimulate egg laying in *unc-13* and PLCβ mutants. These results support a model where serotonin signaling through Gα_q_, PLCβ, and UNC-13 promote neurotransmitter release, and that serotonin also signals through Gα_q_, Trio RhoGEF, and an unidentified, PMA-responsive effector to promote postsynaptic muscle excitability. Thus, the same neuromodulator serotonin can signal in distinct cells and effector pathways to coordinate activation of a motor behavior circuit.

## Introduction

Neurons communicate in circuits via synaptic transmission to initiate, sustain, and terminate behaviors. During neurotransmission, both synaptic vesicles and dense-core vesicles fuse with the presynaptic membrane, releasing neurotransmitters and neuropeptides that activate postsynaptic ion channels and G protein coupled receptors (GPCRs) (Betke et al., 2012; Geppetti et al., 2015). While much has been learned about neurotransmitter signaling pathways through ionotropic receptors, the diversity of GPCRs and their signaling pathways has complicated our understanding of how their signaling exerts changes on cell excitability and behavior. The G protein, Gα_q_, is one of the major G proteins expressed in all excitable cells (Offermanns, 2001; Simon et al., 1991; Wilkie et al., 1992). Activated Gα_q_ signals through PIP_2_-specific Phospholipase-Cβ (PLCβ) to generate the second messengers, inositol 1,4,5 trisphosphate (IP_3_) and 1,2-diacylglycerol (DAG). IP_3_ activates the IP_3_ receptor to release Ca^2+^ from intracellular stores and activate downstream kinases, lipases, and ion channels (Berridge et al., 2000; Huang, 1989; Li et al., 2014; Mujica and Gonzalez, 2011). The membrane lipid DAG has been shown to recruit and activate numerous effector proteins including UNC-13 and Protein Kinase C (Ananthanarayanan et al., 2003; Brose and Rosenmund, 2002; Lou et al., 2008; Maruyama and Brenner, 1991; Rozengurt et al., 1997; Silinsky and Searl, 2003; Thore et al., 2005), but whether these or other identified DAG targets function to transduce all forms of Gα_q_ signaling *in vivo* remains an open question.

Genetic studies in the nematode worm *C. elegans* show that Gα_q_ signaling through both PLCβ and Trio promotes neurotransmitter and neuropeptide transmission. In *C. elegans unc-73* gene encodes at least 8 isoforms of Trio, which has both Rac and Rho GTPase exchange factor (GEF) DH/PH domains (Steven et al., 2005; Williams et al., 2007). *unc-73* mutations that specifically affect the second Rho activating DH/PH GEF domain of Trio (Trio RhoGEF) disrupt locomotion, feeding, and egg-laying behaviors (Williams et al., 2007), without causing the axon pathfinding and neurodevelopmental defects observed in animals bearing *unc-73* Trio RacGEF mutations that affect the first, Rac activating DH/PH GEF domain (Steven et al., 1998). In worms, Gα_q_ knockouts are lethal while PLCβ or Trio RhoGEF single knockouts show defects in neurotransmission that disrupt locomotion, feeding, egg-laying and other behaviors, resembling Gα_q_ loss-of-function mutants (Bastiani et al., 2003; Brundage et al., 1996; Hajdu-Cronin et al., 1999; Lackner et al., 1999). Worms bearing mutations that disrupt both PLCβ and Trio RhoGEF phenocopy the larval arrest phenotype of Gα_q_ null mutants, consistent with these two effectors relaying most or all of the relevant Gα_q_ signaling (Williams et al., 2007). Genetic and biochemical studies showed that Gα_q_ binding to and activation of the Trio RhoGEF domain to promote Rho signaling is conserved in mammals (Chhatriwala et al., 2007; Rojas et al., 2007), however, it remains unclear how PLCβ and Rho signaling promotes neurotransmitter and neuropeptide release *in vivo*. The larval lethality of Gα_q_ null mutants can be rescued by the DAG-mimetic phorbol ester, PMA (Reynolds et al., 2005), and PMA can also rescue the egg-laying defects of PLCβ, Trio RhoGEF double mutants (Williams et al., 2007). These results suggest Gα_q_ signaling through both PLCβ and Trio may ultimately converge to regulate DAG levels and the activation of downstream effectors. Both PLCβ and Trio RhoGEF promote acetylcholine (ACh) release from motor neurons that control locomotion (Lackner et al., 1999; Miller et al., 1999; Williams et al., 2007), although mutations in Trio RhoGEF cause behavior defects more aligned with a function in dense core vesicle release (Hu et al., 2011). Gα_q_, Trio, and PLCβ are expressed in the nervous system and in muscles (Bastiani et al., 2003; Lackner et al., 1999; Steven et al., 1998; Steven et al., 2005; Taylor et al., 2021). While re-expression of PLCβ in motor neurons (Lackner et al., 1999) or Trio in all neurons (Williams et al., 2007) rescues the locomotion behavior defects of their mutants, prior work has shown that Gα_q_ has additional functions to promote egg laying in muscles (Bastiani et al., 2003) where Trio RhoGEF is also expressed (Steven et al., 2005). Loss of PLCβ fails to suppress the hyperactive egg-laying phenotypes of Gα_q_ gain-of-function mutants, consistent with Gα_q_ signaling through other effectors like Trio RhoGEF to regulate egg laying (Bastiani et al., 2003). Indeed, *unc-73* RhoGEF mutations strongly suppress the hyperactive egg-laying behavior phenotypes of Gα_o_ mutants unable to inhibit Gα_q_ signaling (Williams et al., 2007).

Using genetics, optogenetics, pharmacology, and Ca^2+^ imaging techniques we have investigated how Gα_q_ and its two effector pathways regulate egg-laying circuit activity and behavior. We find that PLCβ functions in the neurons while Trio RhoGEF signals in both neurons and muscles to promote egg laying. Loss of each of these effectors imparts specific defects in egg-laying behavior that indicate these proteins function in distinct cells to promote egg-laying circuit activity and behavior. Many of these defects can be rescued in part by treatment with phorbol esters that mimic DAG production. Thus, despite Gα_q_ signaling through independent PLCβ and Trio RhoGEF pathways, these effectors may ultimately converge to increase DAG levels which promote egg-laying behavior.

## MATERIALS AND METHODS

### Strains

*C. elegans* worms were maintained at 20 °C on Nematode Growth Medium (NGM) agar plates with *E. coli* OP50 as a source of food as described previously (Brenner, 1974). All behavior assays and fluorescence imaging experiments were performed with age-matched adult hermaphrodites aged 24-36 h after the late L4 stage. Strains used in this study are listed in Table 1.

**Table 1:** *C. elegans* strains used in this study.

| Strains | Genotype | Feature | Source |
| --- | --- | --- | --- |
| N2 | Wild type | Bristol wild-type strain | (Brenner, 1974) |
| MT1434 | *egl-30(n686) I* | $\text{G}\alpha_q$ loss-of-function mutant, Egg-laying defective | (Trent et al., 1983) |
| DA823 | *egl-30(ad805) I* | $\text{G}\alpha_q$ strong loss-of-function mutant, Egg-laying defective | (Brundage et al., 1996; Mendel et al., 1995) |
| KG1278 | *unc-73(ce362) I* | Trio RhoGEF loss-of-function mutant, Egg-laying defective | (Williams et al., 2007) |
| KG1397 | *unc-73(ev802) I* | Trio RhoGEF deletion mutant, Egg-laying defective | (Williams et al., 2007) |
| JT47 | *egl-8(sa47) V* | $\text{PLC}\beta$ null, Egg-laying defective | (Thomas, 1990) |
| MT1083 | *egl-8(n488) V* | $\text{PLC}\beta$ null, Egg-laying defective | (Trent et al., 1983) |
| CB6614 | *egl-8(e2917) V* | $\text{PLC}\beta$ null, Egg-laying defective | (Yook and Hodgkin, 2007) |
| RM1221 | *egl-8(md1971) V* | $\text{PLC}\beta$ null, Egg-laying defective | (Miller et al., 1999) |
| JIP2081 | *egl-23(bln360[n601]) IV; lin-15AB(n765) vsls164 lite-1(ce314) X* | Egg-laying defective | (Thomas, 1990) and this study |
| MT1212 | *egl-19(n582) IV* | Egg-laying defective | (Trent et al., 1983) |
| MT1444 | *egl-2(n693) V* | Egg-laying defective | (Trent et al., 1983) |
| LX1226 | *eat-16(tm761) I* | G$\alpha_q$ RGS null, Hyperactive egg laying | (Porter and Koelle, 2010) |
| CG21 | *egl-30(tg26) I; him-5(e1490) V* | G$\alpha_q$ gain-of-function mutant, Hyperactive egg-laying | (Garcia et al., 2001) |
| LX1832 | *lite-1(ce314) lin-15(n765ts) X* | Used for transgenic line creation | (Gürel et al., 2012) |
| LX1286 | *egl-8(sa47) I; lin-15(n765ts) X* | PLCβ null mutant, Egg-laying defective; multi-vulva | This study |
| LX1674 | *egl-8(sa47) I; lin-15(n765ts) X; vsEx679* | PLCβ null, Egg-laying defective; non-Muv, expresses GFP in neurons from *rgs-1* promoter | This study |
| LX1675 | *egl-8(sa47) I; lin-15(n765ts) X; vsEx680* | PLCβ null, Egg laying defective; non-Muv, expresses PLCβ and GFP in neurons from *rgs-1* promoter | This study |
| MIA26 | *egl-1(n986dm) V* | Lacks HSNs | (Ravi et al., 2018a) |
| LX1918 | *vsls164 lite-1(ce314) lin-15(n765ts) X* | Expresses GCaMP5 and mCherry in the vulval muscles | (Li et al., 2013) |
| MIA109 | *egl-8(sa47) V; vsls164 lite-1(ce314), lin-15(n765ts) X* | Expresses GCaMP5 and mCherry in the vulval muscles of *egl-8(sa47)* mutant | This study |
| MIA139 | *egl-30(n686) I; vsls164 lite-1(ce314) lin-15(n765ts) X* | Expresses GCaMP5 and mCherry in the vulval muscles of *egl-30(n686)* | This study |
| MIA140 | *egl-30(ad805) I; vsls164 lite-1(ce314), lin-15(n765ts) X* | Expresses GCaMP5 and mCherry in the vulval muscles of *egl-30(ad805)* | This study |
| MIA141 | *unc-73(ce362) I; vsls164, lite-1(ce314) lin-15(n765ts) X* | Expresses GCaMP5 and mCherry in the vulval muscles of *unc-73(ce362)* | This study |
| MIA286 | *egl-30(tg26) I; vsls164, lite-1(ce314) lin-15(n765ts) X* | Expresses GCaMP5 and mCherry in the vulval muscles of *egl-30(tg26)* | This study |
| MIA287 | *eat-16(tm761) I; vsls164 lite-1(ce314) lin-15(n765ts) X* | Expresses GCaMP5 and mCherry in the vulval muscles of *eat-16(tm761)* | This study |
| MIA288 | *egl-8(n488) V; vsls164 lite-1 (ce314) lin-15(n765ts) X* | Expresses GCaMP5 and mCherry in the vulval muscles of *egl-8(n488)* | This study |
| MIA296 | *dgk-1(nu62) lite-1(ce314) lin-15(n765ts) X* | Expresses GCaMP5 and mCherry in the vulval muscles of *dgk-1(nu62)* | This study |
| MIA372 | *unc-73(ce362) I; keyEx64* | Expresses GFP in the neurons and mCherry in the muscles of *unc-73(ce362)* | This study |
| MIA373 | *unc-73(ce362) I; keyEx65* | Expresses GFP and Trio RhoGEF-E cDNA in the neurons, and mCherry and Trio RhoGEF-E cDNA in the muscles of *unc-73(ce362)* | This study |
| MIA374 | *unc-73(ce362) I; keyEx66* | Expresses GFP in neurons | This study |
| MIA375 | *unc-73(ce362) I; keyEx67* | Expresses GFP and Trio RhoGEF-E cDNA in neurons | This study |
| MIA376 | *unc-73(ce362) I; keyEx68* | Expresses mCherry in muscles | This study |
| MIA377 | *unc-73(ce362) I; keyEx69* | Expresses mCherry and Trio RhoGEF-E cDNA in muscles | This study |
| LX1836 | *wzls30 IV; lite-1(ce314) lin-15(n765ts) X* | Expresses Channelrhodopsin-2 (ChR2) in HSNs from the *egl-6* promoter | (Collins et al., 2016) |
| MIA229 | *keyls48; lite-1(ce314) lin-15(n765ts) X* | Expresses ChR2 in vulval muscles from the *ceh-24* promoter | (Kopchock et al., 2021) |
| MIA300 | *egl-30(ad805) I; wzls30 IV; lite-1(ce314) X* | Expresses ChR2 in HSN of *egl-30(ad805)* mutant | This study |
| MIA301 | *egl-30(ad805) I; keyls48; lite-1(ce314) X* | Expresses ChR2 in vulval muscles of *egl-30(ad805)* mutant | This study |
| MIA304 | *wzls30 IV; egl-8(n488) V; lite-1(ce314) X* | Expresses ChR2 in HSNs of *egl-8(n488)* mutant | This study |
| MIA305 | *egl-8(n488) V; keyls48; lite-1(ce314) X* | Expresses ChR2 in vulval muscles of *egl-8(n488)* mutant | This study |
| MIA308 | *wzls30 IV; egl-8(sa47) V; lite-1(ce314) X* | Express ChR2 in HSNs of *egl-8(sa47)* mutant | This study |
| MIA309 | *egl-8(sa47) V; keyls48; lite-1(ce314) X* | Expresses ChR2 in vulval muscles of *egl-8(sa47)* mutant | This study |
| MIA247 | *unc-73(ce362) I; wzls30 IV; lite-1(ce314) X* | Expresses ChR2 in HSN of *unc-73(ce362)* mutant | This study 835<br>836 |
| MIA248 | *unc-73(ce362) I; keyls48; lite-1(ce314) X* | Expresses ChR2 in vulval muscles of *unc-73(ce362)* mutant | This study 837 |
| LX1615 | *vsSi3[Punc-103e::unc-103(e1597dm)-GFP::unc-103, cb-unc-119(+)] II* | Expresses gain-of-function A331T UNC-103 ERG channel in the vulval muscles from the *unc-103e* promoter | This study |
| MT7929 | *unc-13(e51) I* | UNC-13 loss-of-function | (Brenner, 1974) |
| EG9631 | *unc-13(s69) I* | UNC-13 null | (Rose and Baillie, 1980) |
| IK105 | *pkc-1(nj1) V* | nPKCε null | (Okochi et al., 2005) |
| IK130 | *pkc-1(nj3) V* | nPKCε null | (Okochi et al., 2005) |
| RB781 | *pkc-1(ok563) V* | nPKCε null | (Consortium, 2012) |
| VC127 | *pkc-2(ok328) X* | cPKC$\alpha/\beta$ null | (Consortium, 2012) |
| MJ500 | *tpa-1(k501) IV* | nPKC$\delta/\theta$ null | (Tabuse et al., 1989) |
| MJ563 | *tpa-1(k530) IV* | nPKC$\delta/\theta$ null | (Tabuse et al., 1989) |

### Molecular biology and transgenes

#### Vulval muscle GCaMP5 strains

Vulval muscle Ca^2+^ activity was recorded using GCaMP5G (Akerboom et al., 2013) which was expressed along with mCherry from the *unc-103e* promoter (Collins and Koelle, 2013), as previously described (Collins et al., 2016; Ravi et al., 2018a). The wild-type reporter strain, LX1918 *vsIs164 [unc-103e::GCaMP5::unc-54 3’UTR + unc-103e::mCherry::unc-54 3’UTR + lin-15(+)] lite-1(ce314) lin-15(n765ts) X* was described previously (Collins et al., 2016). LX1918 males were crossed separately into DA823 *egl-30(ad805) I*, MT1434 *egl-30(n686) I*, JT47 *egl-8(sa47) V*, MT1083 *egl-8(n488) V*, KG1278 *unc-73(ce362) I*, LX1226 *eat-16(tm761) I*, CG21 *egl-30(tg26) I*, *him-5(e1490) V*, or KP1097 *dgk-1(nu62) X* hermaphrodites to generate MIA140 *egl-30(ad805) I; vsIs164 lite-1(ce314) lin-15(n765ts) X*, MIA139 *egl-30(n686) I; vsIs164 lite-1(ce314) lin-15(n765ts) X*, MIA109 *egl-8(sa47) V; vsIs164 lite-1(ce314) lin-15(n765ts) X*, MIA288 *egl-8(n488) V; vsIs164 lite-1(ce314) lin-15(n765ts) X*, MIA141 *unc-73(ce362) I; vsIs164 lite-1(ce314) lin-15(n765ts) X,* MIA287 *eat-16(tm761) I; vsIs164 lite-1(ce314) lin-15(n765ts) X*, MIA286 *egl-30(tg26) I; vsIs164 lite-1(ce314) lin-15(n765ts) X*, and MIA296 *dgk-1(nu62) vsIs164 lite-1(ce314) lin-15(n765ts) X*, respectively. The corresponding gene mutation was confirmed by phenotype, genotype, or both. Presence of *vsIs164* was confirmed observing the mCherry marker, and *lite-1(ce314) X* was confirmed with PCR genotyping. Oligo sequences used for genotyping the corresponding mutations are shown in Table 2.

**Table 2:**
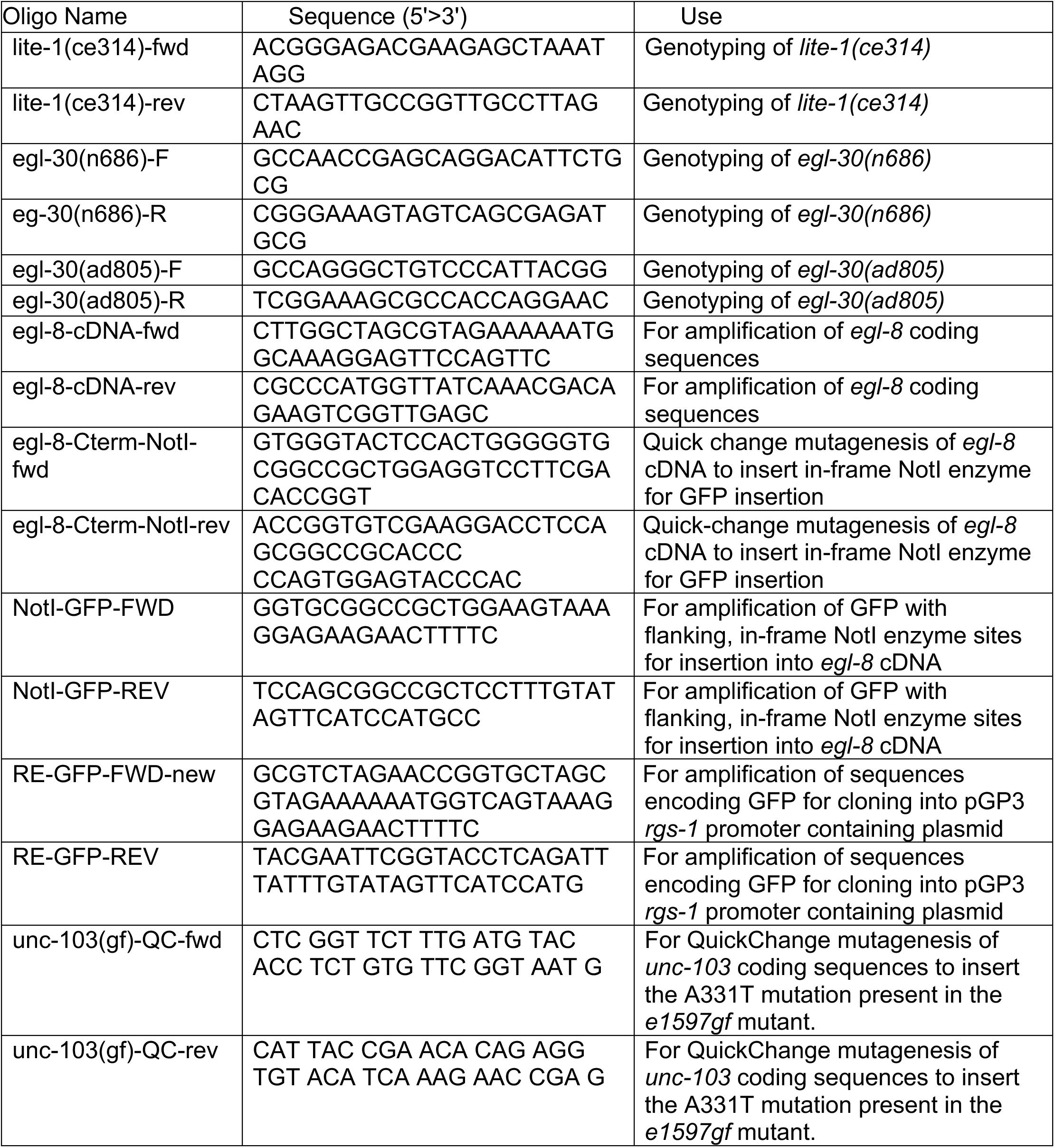
Oligonucleotide sequences used in this study.

### Trio RhoGEF-E transgenes

#### Pan-neuronal expression

The *rab-3* promoter was used to drive expression of GFP alone or with Trio-RhoGEF-E. Briefly, plasmids KG#68 (*rab-3p::GFP*; 15 ng/µl) alone or with KG#281 (*rab-3p:: unc-73e;* 50 ng/µL) were injected into KG1278 *unc-73(ce362) I* (Williams et al., 2007). For behavior experiments, five independent GFP-expressing transgenic lines were used, from which a single transgenic line from each was kept: MIA374 *unc-73(ce362) I*; *keyEx66* (expressing GFP alone) and MIA375 *unc-73(ce362) I*; *keyEx67* (expressing GFP and Trio RhoGEF-E). Plasmids KG#281(*rab-3p::unc-73e*) and KG#68(*rab-3p::GFP*) were kind gifts from Dr. Kenneth Miller.

#### Pan-muscle expression

Plasmid pKMC33 (*rgs-1p::mCherry*) was digested with NheI/KpnI and ligated with similarly digested pPD96.52 (Fire lab *C. elegans* Vector Kit 1999; 1608: L2534, Addgene) to generate pKMC166 (*myo-3p::mCherry*). Plasmid KG#281 (*rab-3p::unc-73e*) was digested with NheI and KpnI, and the insert was ligated into similarly digested pKMC166 to generate pPD3 (*myo-3p::unc-73e*). pKMC166 (15 ng/µL) alone or with pPD3 (50 ng/µL) was injected into KG1278 *unc-73(ce362) I* mutants. Five independent mCherry-expressing transgenic lines were used for behavior experiments from which a single transgenic line from each was kept: MIA376 *unc-73(ce362) I; keyEx68* (expressing mCherry alone) and MIA377 *unc-73(ce362) I*; *keyEx69* (expressing mCherry *+* Trio RhoGEF-E).

#### Neuron and muscle co-expression

Plasmids KG#68 (15 ng/µl; pan-neuronal GFP) or pKMC166 (15 ng/µl; pan-muscle mCherry) alone or with KG#281 (50ng/µl; pan-neuronal *unc-73e*) and pPD3 (50ng/µL; pan-muscle *unc-73e*) were injected into KG1278 *unc-73(ce362) I*, generating five independent mCherry(+), GFP(+) transgenic lines for behavior experiments from which a single transgenic line from each was kept: MIA372 *unc-73(ce362); keyEx64* expressing mCherry (muscles) and GFP (neurons) only and MIA373 *unc-73(ce362); keyEx65* expressing GFP (neurons), mCherry (muscles) and TrioRhoGEF-E (both neurons and muscles).

### PLCβ Transgenes

To generate a control plasmid expressing GFP in neurons, GFP coding sequences were amplified from pJM60 (Moresco and Koelle, 2004) using oligonucleotides RE-GFP-FWD / -REV, digested with NheI / KpnI, and ligated into similarly digested pGP3 bearing the *rgs-1* promoter (Dong et al., 2000), generating pKMC78. An *egl-8* cDNA was used to generate and express a functional GFP fusion protein in neurons. Briefly, oligonucleotides egl-8-cDNA-fwd / -rev were used to amplified *egl-8* coding sequences from a plasmid bearing an *egl-8* cDNA provided by Dr. Kenneth Miller (pKP309). This amplicon was digested with NheI / NcoI and ligated into a similarly digested pPD49.26 plasmid, generating pKMC193. Quickchange mutagenesis with oligonucleotides egl-8-Cterm-NotI-fwd / -rev were used to insert an in-frame NotI site near the 3’ end of the *egl-8* cDNA in a divergent region of the coding sequence, generating plasmid pKMC194. Coding sequences for *egl-8* bearing this NotI site were then moved to pKMC78 by digestion of pKMC194 with NheI/NcoI followed by ligation into a similarly digested pKMC78, generating pKMC195. Oligonucleotides NotI-GFP-FWD / -REV were used to amplified GFP coding sequences from pKMC78, digested with NotI, and ligated into a similarly digested pKMC195, generating pKMC196. A strain bearing the *egl-8(sa47)* mutation was generously provided by Dr. Joshua Kaplan and backcrossed four times to N2 wild-type animals to generate LX1225 *egl-8(sa47) V.* MT8189 *lin-15(n765ts)* males were mated to LX1225 to generate LX1287 *egl-8(sa47) V; lin-15(n765ts) X* hermaphrodites that were kept at 15°C prior to injection. Plasmids expressing GFP alone (pKMC78; 5 ng/µl) or *egl-8* CDNA fused to GFP (pKMC196; 5 ng/µl) from the *rgs-1* promoter were injected along with pL15EK (50 ng/µl) into LX1287 hermaphrodites. For behavior experiments, five independent GFP-expressing lines were used from which a single transgenic line (*vsEx679* [GFP] and *vsEx680* [EGL-8::GFP], respectively) was kept. We noted that transgenic expression of GFP alone did cause a modest, but significant reduction of egg accumulation compared to LX1225 *egl-8(sa47) V* mutant animals. This effect appeared to be specific for egg accumulation, as these same *egl-8(sa47)* GFP-only expressing transgenic lines showed similar resistance to 1 mM aldicarb as LX1225 *egl-8(sa47) V* (0 ±2.1% vs. 9 ±5.9% of animals paralyzed at 4 h, respectively). In contrast, 48 ± 7.9% of transgenic *egl-8(sa47)* animals expressing EGL-8::GFP were paralyzed at 4 h, not significantly different to wild-type N2 animals (51 ±9.6% of animals paralyzed at 4 h).

### Vulval muscle Channelrhodopsin-2 strains

N2 males were crossed into MIA229 *keyIs48 [ceh-24p::ChR2::unc-54 3’UTR + lin-15(+)], lite-1(ce314), lin-15(n765ts) X* (Kopchock et al., 2021) to produce F1 heterozygous males, which then were crossed separately into MIA211 *unc-73(ce362) I; lite-1(ce314) lin-15(n765ts) X,* MIA299 *egl-30(ad805) I; lite-1(ce314) lin-15(n765ts) X*, MIA303 *egl-8(n488) V; lite-1(ce314) lin-15(n765ts) X*, or MIA307 *egl-8(sa47) V; lite-1(ce314) lin-15(n765ts) X* hermaphrodites to generate vulval muscle specific Channelrhodopsin-2 (ChR2) expressing transgenic lines MIA248 *unc-73(ce362) I; keyIs48; lite-1(ce314) X*, MIA301 *egl-30(ad805) I; keyIs48; lite-1(ce314) X*, MIA305 *egl-8(n488) V; keyIs48; lite-1(ce314) X*, and MIA309 *egl-8(sa47) V; keyIs48; lite-1(ce314) X*, respectively. The presence of *lite-1(ce362)* was confirmed by genotyping as above, and the presence of the ChR2 transgene was confirmed by rescue of the *lin-15(n765ts)* multi-vulva (Muv) phenotype.

### HSN Channelrhodopsin-2 strains

ChR2 was expressed in the HSNs from the *egl-6* promoter via an integrated *wzIs30* transgene (Emtage et al., 2012). This transgene was crossed into Gα_q_ signaling mutants as follows. N2 males were crossed into LX1836 *wzIs30 IV; lite-1(ce314) lin-15(n765ts) X* to generate heterozygous F1 males, which were then crossed separately into MIA211 *unc-73(ce362) I; lite-1(ce314) lin-15(n765ts) X,* MIA299 *egl-30(ad805) I; lite-1(ce314) lin-15(n765ts) X*, MIA303 *egl-8(n488) V; lite-1(ce314) lin-15(n765ts) X,* or MIA307 *egl-8(sa47) V; lite-1(ce314) lin-15(n765ts) X* hermaphrodites to generate MIA247 *unc-73(ce362) I; wzIs30 IV; lite-1(ce314) X,* MIA300 *egl-30(ad805) I; wzIs30 IV; lite-1(ce314) X*, MIA304 *wzIs30 IV; egl-8(n488)V; lite-1(ce314) X*, and MIA308 *wzIs30 IV; egl-8(sa47) V; lite-1(ce314) X*, respectively. The presence of *lite-1(ce362)* was confirmed by PCR genotyping, and the *wzIs30* transgene was confirmed by rescue of the *lin-15(n765ts)* Muv phenotype.

### Other strains

A single-copy MosSCI knock-in strain expressing *unc-103* bearing the *e1597dm* gain-of-function mutation was constructed as described (Collins and Koelle, 2013). Briefly, plasmid pKMC179 bearing *unc-103* coding sequences behind the *unc-103e* promoter/enhancer was mutagenized by QuickChange using primers unc-103(gf)-QC-fwd and unc-103(gf)-QC-rev to generate pKMC183. Digestion with BsrGI and Sanger sequencing confirmed the mutagenesis was successful. *Unc-103(e1597gf)* coding sequences were then PCR-amplified from pKMC183 using Phusion polymerase (NEB), digested with NheI/MluI enzymes, and then ligated into pKMC176, a *ttTi5606* site MosSCI donor plasmid, generating pKMC185. pKMC185 was then injected at 50 ng/µl along with plasmids expressing Mos1 transposase and mCherry co-injection markers into EG4322 *ttTi5605 II; unc-119(ed9) III*, as described (Frokjaer-Jensen et al., 2008), generating LX1565 *vsSi3[Punc-103e::unc-103e(e1597dm)-GFP, cb-unc-119(+)] II*; *unc-119(ed3) III* which had a strong Egl phenotype but markedly reduced Unc phenotype compared to the reference CB1597 *unc-103(e1597dm)* strain. LX1565 was then outcrossed to N2 four times to generate LX1615 *vsSi3[Punc-103e::unc-103e(e1597dm)-GFP, cb-unc-119(+)] II*.

### Behavior assays

Quantification of egg accumulation was performed as described (Chase and Koelle, 2004). Staged adults were obtained by picking late L4 animals and culturing them 24-30 h at 20 °C. Each animal was placed in 7 µL of 20% hypochlorite (bleach) solution and eggs were counted after animals had dissolved. Numbers of eggs and any internally hatched L1 animals were combined.

### Pharmacological assays

Egg laying in response to exogenous serotonin was performed as described (Banerjee et al., 2017; Kopchock et al., 2021). Individual staged adult animals were placed in 100 μl of either M9 buffer alone, or M9 containing 18.5 mM serotonin (creatinine sulfate monohydrate salt, Sigma-Aldrich #H7752) or M9 containing 10 µM PMA (Phorbol-12-myristate-13-acetate, Calbiochem #524400) in a 96-well microtiter dish. After 1 hour, the number of released eggs and L1 larvae in each well were counted. Since egg-laying defective animals sometimes release one or two eggs/L1 larvae when they are first picked into the well in response to mechanical stimulation, animals were only recorded as responding if they laid 3 or more progeny. For Ca^2+^ imaging, NGM plates containing either PMA or ethanol solvent were prepared as described (Reynolds et al., 2005). Age-matched adult worms from each genotype were placed on separate PMA or control NGM plates at room temperature for 1.5 hours. An agar chunk was then placed between two glass coverslips for Ca^2+^ activity recording as described (Ravi et al., 2018b). The unused plates were kept at 4 °C for future use.

### Optogenetic assay

All-*trans* retinal (Sigma Aldrich, R2500) was resuspended in ethanol (100%) to make 100 mM solution and added to a warmed culture of OP50 bacteria grown in B Broth media to a final concentration of 0.4 mM. Individual NGM agar plates were seeded with 200 μl of freshly prepared +ATR food and were grown in the dark for ∼24 hours prior to use. In all photo-stimulation experiments, a set of control animals were grown in the absence of ATR. Animals were imaged at 4x magnification on a Leica M165FC stereomicroscope and illuminated with 3.3 mW/cm^2^ of ∼470 ± 20 nm blue light produced using a EL6000 metal halide light source and a GFP excitation/emission filter set. The 30 s on/off sequence was programmed and controlled using a Doric Optogenetics TTL Pulse Generator (OTPG-4, Version 3.3) triggering a SHB1 series shutter controller (ThorLabs; 170712-1).

### Microscopy

#### Ratiometric Ca^2+^ imaging

Vulval muscle Ca^2+^ activity was performed in freely behaving adult animals at 24-30 h past the late L4 larval stage, as described previously (Collins et al., 2016; Collins and Koelle, 2013; Ravi et al., 2018a). Worms co-expressing GCaMP5G and mCherry under the *unc-103e* promoter transgene *vsIs164* were mounted beneath the chunk of agar over the glass coverslip, and reporter fluorescence was recorded through an 20X Apochromatic objective (0.8 NA) mounted on an inverted Zeiss Axio Observer.Z1. A Colibri.2 LED illumination system was used to excite GCaMP5 at 470 nm and mCherry at 590 nm for 10 msec every 50 msec. GFP and mCherry fluorescence emission channels were separated using a Hamamatsu W-VIEW Gemini image splitter and recorded simultaneously for 10 min with an ORCA-Flash 4.0 V2 sCMOS camera at 256 / 256-pixel resolution (4×4 binning) at 16-bit depth. A motorized stage was manually controlled using a joystick to maintain the freely behaving animal in the field of view. For experiments without treatment of PMA or vehicle control, animals were recorded until each entered into an egg-laying active state. The recording was then cropped to a 10 minute (12,000 frame) two channel image sequence, centered on the first egg-laying event observed, for subsequent ratiometric analysis. The egg-laying active state was operationally defined as starting 1 minute before the first egg-laying event and ending 1 minute after the last egg-laying event observed in the 10 min recording. For egg-laying defective mutants like *egl-30(ad805)* and *unc-73(ce362)* that lay essentially no eggs, the 10 min extraction was not centered on any specific behavioral feature. For drug and vehicle control assays, the 10 min recording period started immediately, whether or not animals laid eggs or were seen to enter into an active state. Image sequences were exported to Volocity software (Quorum Technologies Inc.) for segmentation and ratiometric analysis. Ca^2+^ transient peaks from ratio traces were detected using a custom MATLAB script, as described (Ravi et al., 2018b).

### Experimental design and statistical analysis

Sample sizes for behavioral assays followed previous studies (Chase and Koelle, 2004; Collins et al., 2016). Statistical analysis was performed using Prism v.8 or v.9 (GraphPad). Ca^2+^ transient peak amplitudes, widths, and inter-transient intervals were pooled from multiple animals (typically ≥10 animals per genotype). All statistical tests were corrected for multiple comparisons (Bonferroni for one-way ANOVA or Fisher’s exact tests; Dunn’s correction for Kruskal–Wallis tests). Each figure legend indicates individual p-values with p<0.05 being considered significant.

### Data Availability statement

All the data, reagents, and strains used in this study are available from the corresponding author upon request.

## Results

### Trio RhoGEF acts in both neurons and muscles to drive egg-laying behavior

Prior studies have shown that Gα_q_ signaling (Fig. 1A) through PLCβ and Trio RhoGEF promotes neurotransmitter release and locomotion (Brundage et al., 1996; Miller et al., 1999; Williams et al., 2007). How Gα_q_ signaling regulates other *C. elegans* behavior circuits is less well established. To address this uncertainty, we chose to examine the neural circuit driving egg-laying behavior. Egg-laying behavior in *C. elegans* is regulated by a small motor circuit with defined neurons and muscle connectivity (Cook et al., 2019; White et al., 1986). The HSNs are serotonergic command motor neurons (Fig. 1B) that initiate the egg-laying active state and promote the excitability of the vm1 and vm2 egg-laying vulval muscles (Collins et al., 2016; Emtage et al., 2012; Waggoner et al., 1998). Innervating ventral cord motor neurons release acetylcholine to regulate muscle contraction (Bany et al., 2003; Kim et al., 2001; Kopchock et al., 2021; Waggoner et al., 2000). We first examined the steady-state accumulation of eggs in the uterus as a proxy for changes in egg-laying circuit activity and behavior. As previously shown, animals bearing mutations in the EAT-16 RGS protein, which inhibits Gα_q_ signaling (Hajdu-Cronin et al., 1999), or gain-of-function mutations in Gα_q_ itself, (Bastiani et al., 2003; Doi and Iwasaki, 2002) showed a significant increase in egg laying resulting in a significant reduction in egg accumulation compared to the ∼15 ± 1.4 embryos retained in wild-type animals (Fig. 1C-E; Fig. 1I). Conversely, animals bearing mutations which reduce Gα_q_ signaling showed the opposite phenotype. Animals bearing an early nonsense mutation predicted to be a PLCβ null mutant*, egl-8(sa47),* (Lackner et al., 1999; Williams et al., 2007) accumulated an average of 21 eggs, showing a significant increase in egg retention (Fig. 1G). Animals bearing a missense mutation in the RhoGEF domain of Trio, *unc-73(ce362)* (Williams et al., 2007), showed an even stronger egg-laying behavior defect, accumulating more than 30 eggs in the uterus, closely resembling animals bearing loss-of-function mutations in Gα_q_ itself (Fig. 1F-1I). Together, these results confirm that Gα_q_ and its effectors PLCβ and Trio RhoGEF are required for egg-laying behavior in *C. elegans* and that loss of the Trio RhoGEF branch causes a stronger behavior impairment compared to loss of PLCβ.

**Figure 1:**
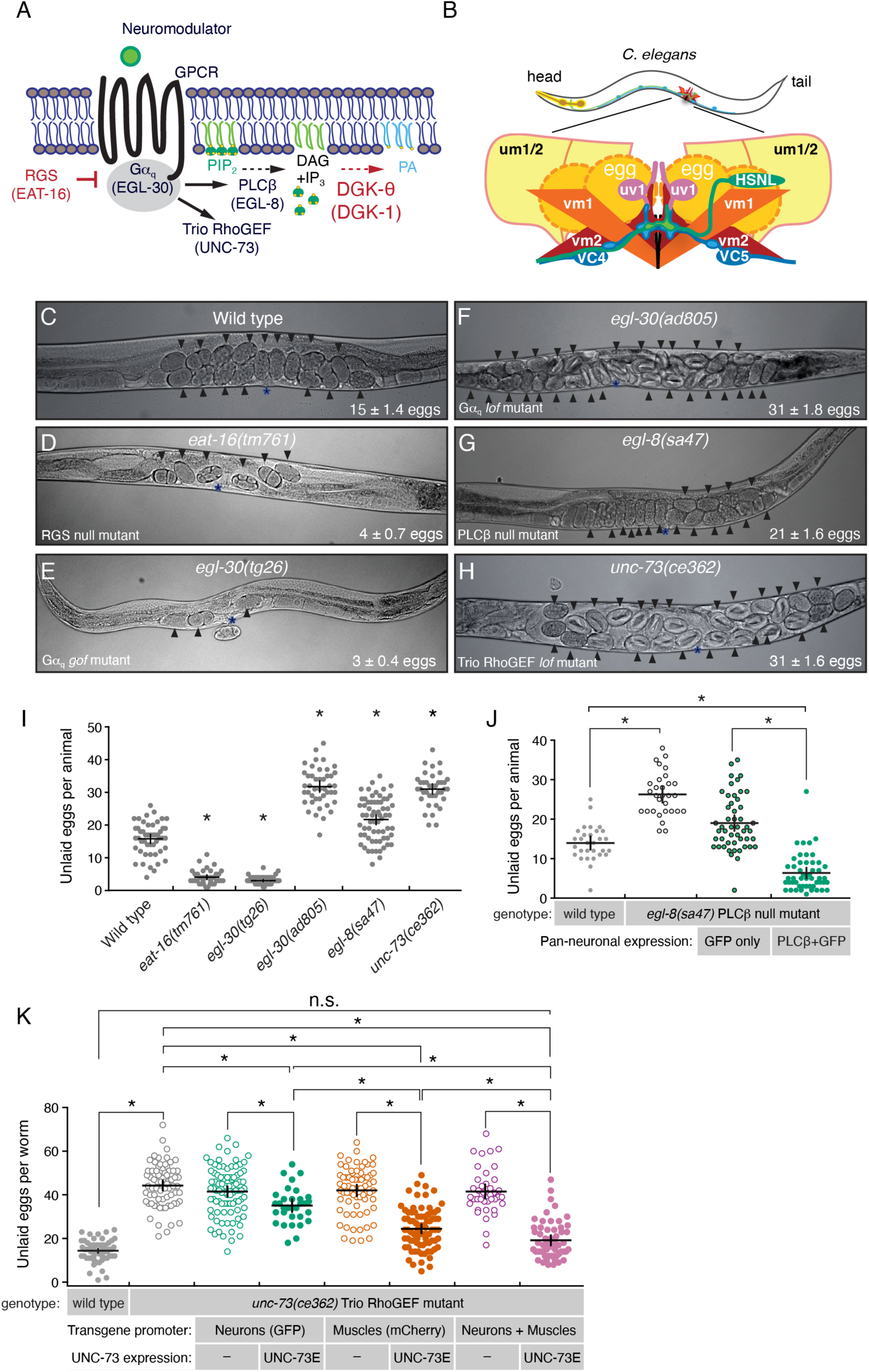
Trio RhoGEF acts in both neurons and muscles to regulate egg-laying behavior. **(A)** Schematics of excitatory (black) and inhibitory (red) Gα_q_ signaling pathway. *C. elegans* gene names are beneath the protein they encode. **(B)** Cartoon of the *C. elegans* egg-laying circuit from a lateral view. Only the left side of the bilaterally symmetric circuit is shown. HSNL, Hermaphrodite Specific Neuron (left); VC4 and VC5 Ventral C neurons; vm1 and vm2 vulval muscles, um1 and um2 uterine muscles; uv1 uterine-vulval neuroendocrine cells. **(C-H)** Bright field images of worms of the indicated genotypes; arrowheads indicate accumulated eggs. Mean number of accumulated eggs ± 95% confidence intervals is also indicated. Position of the vulva is shown with an asterisk (*). **(I)** Scatterplot of egg accumulation in wild-type, *eat-16(tm761)*, *egl-30(tg26)*, *egl-30(ad805), egl-8(sa47), and unc-73(ce362)* mutant animals. Line indicates mean eggs ± 95% confidence intervals. Asterisks indicate p ≤0.0001 (One way ANOVA with Bonferroni’s correction; wild type (n=49); *eat-16(tm761)* (n=36); *egl-30(tg26)* (n=47); *egl-30(ad805)* (n=44); *egl-8(sa47)* (n=65); *unc-73(ce362)* (n=38). **(J)** Transgenic rescue of *egl-8* PLCβ egg-laying defects. Scatterplot of egg accumulation in transgenic animals expressing GFP only or EGL-8/PLCβ fused to GFP expressed from the *rgs-1* promoter in *egl-8(sa47)* mutants (n=50) compared to wild-type (n=30) and *egl-8(sa47)* mutant animals (n=30). Bar indicates mean eggs ±95% confidence intervals. Asterisks indicate p≤0.0001 (One-way ANOVA with Bonferroni’s correction) **(K)** Transgenic rescue of *unc-73* Trio RhoGEF egg-laying defects. Scatterplot of egg accumulation in wild-type (n=60), *unc-73(ce362)* mutants (n=72), and transgenic animals expressing a fluorescent protein with or without Trio/UNC-73E in neurons from the *rab-3* promoter (n≥32) or in muscles from the *myo-3* promoter (n≥69), or in both neurons and muscles (n≥41) in *unc-73(ce362)* mutants. Horizontal line indicates mean accumulated eggs ±95% confidence intervals. Asterisks indicate p≤0.0145; n.s.= not significant (p>0.05; One-way ANOVA with Bonferroni’s correction for multiple comparisons).

Previous work has shown that Gα_q,_ Trio, and PLCβ are expressed in neurons and muscles of the egg-laying circuit (Bastiani et al., 2003; Brundage et al., 1996; Lackner et al., 1999; Miller et al., 1999; Steven et al., 1998; Taylor et al., 2021). To understand where Gα_q_ and its effectors function to regulate egg laying, we used tissue-specific promoters to express cDNAs encoding PLCβ or Trio RhoGEF in either all neurons, in the body wall and egg-laying vulval muscles, or in both neurons and muscles. We found that transgenic expression of PLCβ from the pan-neuronal *rgs-1* promoter (Dong et al., 2000) in PLCβ null mutants was sufficient to rescue their defects in egg laying (Fig. 1J) and acetylcholine (ACh) release as measured by restoration of sensitivity to aldicarb, a cholinesterase inhibitor (see Methods). This suggests PLCβ functions in neurons to regulate egg laying. Previous work has indicated the presence of eight transcript variants of Trio (A, B, C1, C2, D1, D2, E, and F), which are differentially expressed in *C. elegans* (Steven et al., 1998; Steven et al., 2005). Transgenic expression of Trio RhoGEF-E in neurons is sufficient to rescue the locomotion defects of Trio RhoGEF mutants (Williams et al., 2007). To explore whether Trio RhoGEF acts similarly in neurons to promote egg laying, we used the *rab-3* pan-neuronal promoter (Williams et al., 2007) to express Trio RhoGEF-E and measured egg accumulation in these animals. We observed a modest, but significant reduction in the number of eggs retained in Trio RhoGEF mutants (∼36 eggs) compared to control Trio RhoGEF mutant animals (∼42 eggs; Fig. 1K). Transgenic expression of Trio RhoGEF-E in the egg-laying vulval muscles from a muscle-specific promoter showed a greater rescue of egg accumulation (∼25 eggs), and this rescue of egg laying was improved to nearly wild-type levels when Trio RhoGEF-E was expressed in both neurons and muscles (∼19 eggs; Fig. 1K). Although these rescue lines are extrachromosomal arrays and likely expressed at different levels and with different amounts of mosaicism, these results support a general interpretation that *unc-73* functions in both neurons and muscles to promote egg laying. However, because previous work showed that expression of *unc-73* in neurons can rescue locomotion defects (Williams et al., 2007) but not egg laying (Fig. 1K and data not shown), these results indicate that *unc-73* has additional functions in muscle that cannot be bypassed or rescued by expression just in neurons. Together, these results suggest that Gα_q_ effectors PLCβ and Trio RhoGEF function in neurons to regulate egg-laying behavior. Our results also suggest Trio RhoGEF also functions in the postsynaptic vulval muscles for proper regulation of egg laying, a finding consistent with previously results regarding Gα_q_ (Bastiani et al., 2003).

### Serotonin signals through Gα_q_, Trio, and PLCβ to promote egg laying

Previous studies have shown that serotonin released from the HSNs signals through G-protein coupled serotonin receptors expressed on the vulval muscles (Bastiani et al., 2003; Dempsey et al., 2005; Fernandez et al., 2020; Tanis et al., 2008; Xiao et al., 2006). The vulval muscles are also innervated by cholinergic ventral cord motor neurons (Cook et al., 2019; White et al., 1986) whose release of ACh is regulated by serotonin and G protein signaling (Nurrish et al., 1999). To test how serotonin promotes egg laying via Gα_q_, we measured the egg-laying response to serotonin in Gα_q_ signaling mutants. Serotonin promotes egg laying in hypertonic M9 buffer, a condition that normally inhibits egg laying in both wild-type and HSN-deficient *egl-1(n986dm)* mutants, which developmentally lack the HSNs (Fig. 2B). Consistent with previous results (Bastiani et al., 2003; Brundage et al., 1996; Trent et al., 1983), more than 62% of wild-type animals and 57% of HSN-deficient *egl-1(n986dm)* mutant animals laid eggs in response to serotonin compared to only 13% of Gα_q_ mutant animals (Fig. 2B). Serotonin response was similarly and significantly reduced to 23% and 3% in PLCβ and Trio RhoGEF mutant animals, respectively (Fig. 2B). Our results are consistent with the previous data reporting that Gα_q_ loss-of-function mutants and the PLCβ deletion mutant, *egl-8(n488)* do not lay eggs in response to exogenous serotonin (Bastiani et al., 2003; Trent et al., 1983).

**Figure 2:**
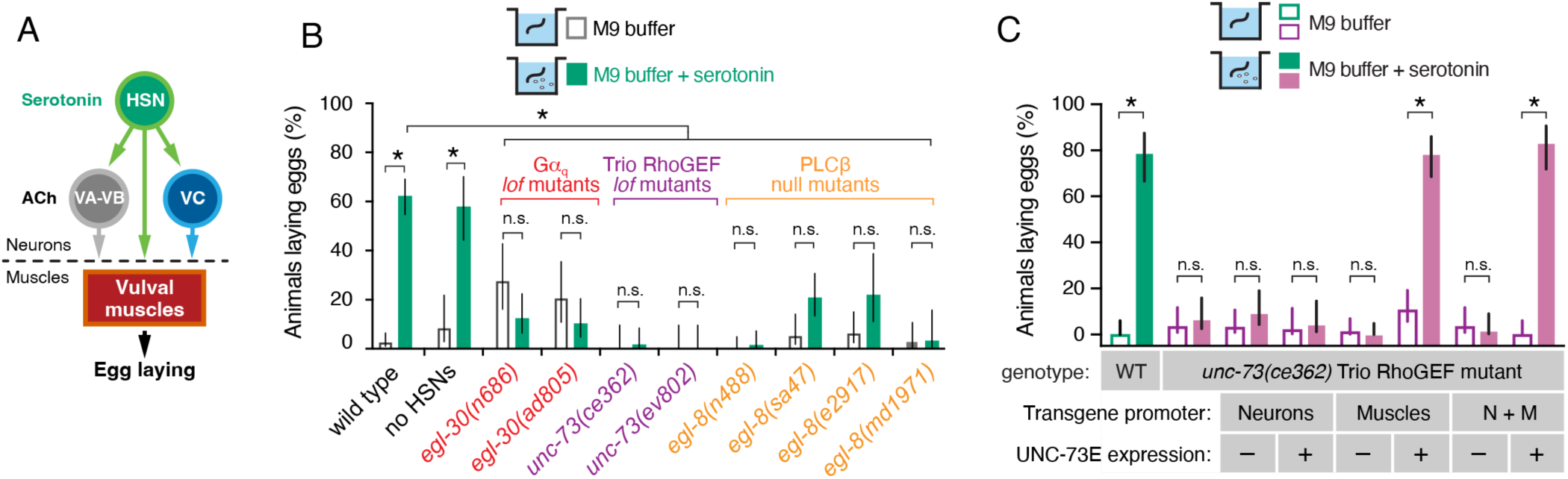
Serotonin signals through Gα_q_, Trio, and PLCβ to promote egg laying. **(A)** Model of serotonergic and acetylcholine (ACh) signaling in the egg-laying circuit. **(B)** Bar plots showing the percentage of animals laying eggs in M9 buffer alone (grey) or M9 +18.5 mM serotonin (green) after 1 hr. Bar indicates mean percent ±95% confidence intervals. Asterisks indicate p<0.0007; n.s.= not significant (p>0.05, Fisher’s exact test with Bonferroni’s correction for multiple comparisons; n>30 animals for each genotype and condition). **(C)** Bar plot showing percent of animals laying eggs in M9 buffer or M9 +18.5 mM serotonin in wild-type or Trio RhoGEF mutant animals expressing nothing or Trio/UNC-73E in neurons, muscles, or both. Bar indicates mean percent ±95% confidence intervals. Asterisks indicate p<0.0007; n.s.= not significant (p>0.05, Fisher’s exact test with Bonferroni’s correction for multiple comparisons; n>30 animals).

To confirm whether PLCβ is required for egg laying in response to serotonin, we tested other PLCβ mutants including *sa47* and *md1971*, both of which carry nonsense mutations predicted to terminate the protein prematurely, and *e2917* in which the coding sequence is disrupted by a Mos1 transposon (Lackner et al., 1999; Miller et al., 1999; Yook and Hodgkin, 2007). All PLCβ mutants tested failed to lay eggs in response to exogenous serotonin after 60 min (Fig. 2B), but only *egl-8(n488)* animals remained resistant to serotonin after 90 minutes (Table 3), with the other PLCβ mutant animals beginning to lay eggs by 90 minutes, consistent with previous observations (Bastiani et al., 2003). Taken together, these results indicate that Gα_q_, PLCβ, and Trio act at least in part outside of HSNs to promote egg laying in response to serotonin. To determine where in the animal the Trio RhoGEF deficiency caused serotonin insensitivity, we measured egg laying in Trio RhoGEF mutant animals re-expressing Trio RhoGEF-E in either neurons, muscles, or both. Transgenic expression of Trio RhoGEF-E in neurons failed to rescue egg laying in response to serotonin (Fig. 2C), but expression of Trio RhoGEF-E in muscles, or in both neurons and muscles, restored egg laying of *unc-73(ce362)* mutant animals (Fig. 2C), suggesting Trio RhoGEF mediates serotonin signaling by acting in the vulval muscles. Together, these results indicate that Gα_q_, PLCβ, and Trio RhoGEF function at least in part outside of the HSNs to drive egg laying in response to serotonin with Trio RhoGEF likely functioning in the muscles.

**Table 3:** Serotonin-induced egg laying in *egl-8* PLCβ mutants.

|  |  | 60 min |  | 90 min |  | P value<br>(Paired t test) |
| --- | --- | --- | --- | --- | --- | --- |
| Genotype | N | Average eggs laid in<br>18.5 mM serotonin | Std.<br>Deviation | Average eggs laid in<br>18.5 mM serotonin | Std.<br>Deviation |  |
| wild type (N2) | 64 | 1.8 | 1.7 | 5.8 | 4.5 | <0.0001 |
| *egl-8(e2917)* | 32 | 0.2 | 0.6 | 1.4 | 2.5 | 0.0091 |
| *egl-8(n488)* | 32 | 0.0 | 0.2 | 0.1 | 0.3 | 0.0831 |
| *egl-8(md1971)* | 32 | 0.3 | 0.8 | 1.3 | 2.1 | 0.0075 |
| *egl-8(sa47)* | 32 | 0.8 | 1.4 | 2.5 | 2.5 | 0.0002 |

### Optogenetic stimulation of the HSNs and vulval muscles suggests cellular specificity of Gα_q_ effectors for egg-laying

Optogenetic stimulation of Channelrhodopsin-2 (ChR2) expressed in either the HSNs (Emtage et al., 2012) or vulval muscles (Kopchock et al., 2021) can drive egg laying. To test whether and how Gα_q_ and its effectors mediate this response, we expressed ChR2 in HSNs in Gα_q_ and effector mutants and measured egg laying during 30s of exposure to blue light. Blue light stimulation of the HSNs drove an average release of ∼3 eggs in wild-type animals, which was reduced to essentially zero in Gα_q_ [(*egl-30(ad805)*] and Trio RhoGEF [*unc-73(ce362)*] mutants (Fig. 3A), consistent to our previous results showing Trio RhoGEF acts in part downstream of the HSNs in the postsynaptic vulval muscles. In contrast, optogenetic stimulation of the HSNs in PLCβ null mutants [*egl-8(sa47)* or *egl-8(n488)*] drove robust egg release, releasing an average of ∼4 and ∼6 embryos respectively in 30 s (Fig. 3A). To test whether the failure of egg laying in Gα_q_ and Trio RhoGEF mutants was a consequence of muscle developmental defects, rather than excitability deficits, we expressed and stimulated ChR2 in the vulval muscles. Blue light exposure drove the release of ∼4 eggs in 30 s in wild-type animals. Both PLCβ [*egl-8(sa47)* and *egl-8(n488)*] and Trio RhoGEF mutants [*unc-73(ce362)*] laid a similar number of eggs as wild-type control animals after blue light stimulation (Fig. 3B). The *egl-30(ad805)* Gα_q_ mutant laid slightly fewer eggs on average (∼3) but this was not significant. Thus, the failure of Gα_q_ and Trio mutants to lay eggs in response to exogenous serotonin or optogenetic stimulation of the HSNs does not arise from some intrinsic defect in vulval muscle contractility, but rather a specific deficiency in muscle excitability. These results are also consistent with prior findings showing rescue of egg-laying behavior defects of Gα_q_, PLCβ, and Trio RhoGEF mutants by exogenous phorbol esters (Lackner et al., 1999; Williams et al., 2007).

**Figure 3:**
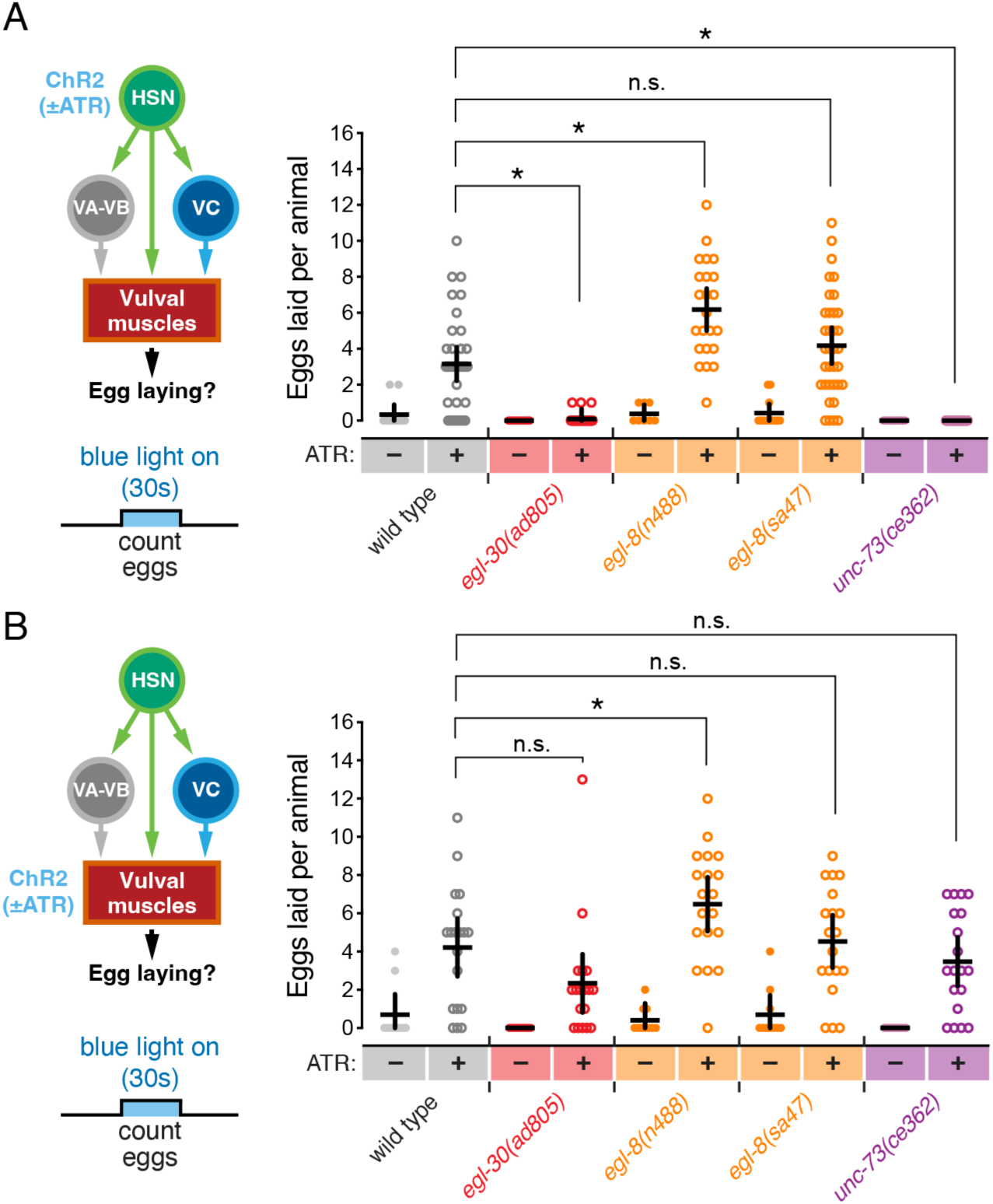
Optogenetic stimulation of the HSNs or vulval muscles reveals distinct cellular specificity of Gα_q_ effectors for egg laying. **(A)** On left, cartoon of the egg-laying circuit and experiment showing blue light activation of HSN for 30 seconds. On right, scatterplot showing eggs laid per worm in presence (+) or absence (-) of all-*trans* retinal (ATR) cofactor during the blue light activation of ChR2 expressed in HSNs of wild-type (grey), *egl-30(ad805)* Gα_q_ strong loss-of-function mutants (red), *egl-8(n488) and egl-8(sa47)* PLCβ mutants (orange), and *unc-73(ce362)* Trio mutant (purple) animals. Line indicates mean eggs laid ±95% confidence intervals. Asterisks indicate p<0.0001; n.s.= not significant (p>0.05, One-way ANOVA with Bonferroni’s correction for multiple comparisons; n>10). (**B)** On the left, cartoon of the egg-laying circuit and experiment showing blue light activation of vulval muscles for 30 seconds(left). On the right, scatter plots of eggs laid per worm in presence (+) or absence (-) of ATR during blue light activation of ChR2 expressed in the vulval muscles of wild type, *egl-30(ad805)* Gα_q_ strong loss-of-function mutants (red), *egl-8(n488) and egl-8(sa47)* PLCβ mutants (orange), and *unc-73(ce362)* Trio mutant (purple) animals. Line indicates mean eggs laid ±95% confidence intervals. Asterisk indicates p≤0.0255; n.s.=not significant, p>0.05 (One-way ANOVA with Bonferroni’s correction for multiple comparisons; n>10).

Collectively, we interpret the previous serotonin experiments and these optogenetic results as showing that Gα_q_ signals through PLCβ in HSN and/or in other neurons to promote release of neurotransmitters that signal through vulval muscle receptors coupled to Gα_q_ and Trio RhoGEF. That HSN optogenetic stimulation, but not exogenous serotonin, stimulates egg laying in PLCβ mutants suggests HSN releases other factors such as NLP-3 neuropeptides (Brewer et al., 2019) which signal to promote egg laying in parallel to serotonin and PLCβ. Our data do not support a model where PLCβ acts only in HSNs as *egl-1(n986dm)* animals lacking HSNs still lay eggs in response to serotonin while *egl-8* PLCβ null mutants do not (Fig. 2B). Together, these results support the conclusion from our rescue experiments (Fig. 1J-1K) that PLCβ and Trio RhoGEF function in distinct cells and through unique mechanisms to promote egg-laying circuit activity and behavior.

### Gα_q_ and Trio RhoGEF are required for vulval muscle Ca^2+^ activity

Egg laying is a two-state behavior where ∼20 minute inactive states are punctuated by ∼2-minute active states with high levels of rhythmic Ca^2+^ transient activity in the egg-laying circuit driving release of 3-5 eggs (Collins et al., 2016; Waggoner et al., 1998; Zhang et al., 2008; Zhang et al., 2010). Loss of Gα_q_ signaling in *egl-30(n686)* animals causes a significant reduction in spontaneous and serotonin-induced vulval muscle Ca^2+^ transients in immobilized animals (Shyn et al., 2003). We therefore tested whether these Gα_q_ dependent Ca^2+^ activity defects were similarly seen in freely behaving animals on solid media and whether they were shared in PLCβ and Trio RhoGEF mutants. We expressed the genetically encoded Ca^2+^ reporter, GCaMP5, along with mCherry in the vulval muscles of mutant animals with either too much or too little Gα_q_ signaling and performed ratiometric imaging as they entered and left the egg-laying active state.

As shown in Figure 4A, the normal two-state pattern of rhythmic vulval muscle Ca^2+^ activity (and egg laying) is lost in animals bearing strong loss-of-function Gα_q_ or Trio RhoGEF mutations. To further quantify these activity defects, we compared vulval muscle Ca^2+^ transient amplitudes and frequencies in wild-type and Gα_q_ signaling mutant animals. In *egl-30(ad805)* Gα_q_ and *unc-73(ce362)* Trio mutants which laid no eggs during the recording period, we failed to see the large amplitude egg-laying Ca^2+^ transients (1.4 ± 0.1 ΔR/R) typically observed in wild-type animals where both the vm1 and vm2 muscles contract or even the smaller amplitude rhythmic ‘twitch’ Ca^2+^ transients (0.4 ± 0.03 ΔR/R) that are localized primarily to the vm1 muscles (Fig. 4A and 4B). As a result, the wild-type frequency of 3.5 ± 1.0 (ΔR/R) Ca^2+^ transients per min was significantly reduced to essentially zero in *egl-30(ad805)* Gα_q_ and *unc-73(ce362)* Trio mutants (Fig. 4C). In contrast, we did not observe a significant reduction in the amplitude or frequency of vulval muscle Ca^2+^ transients in *egl-30(n686)* Gα_q_ weak loss-of-function mutants or *egl-8(sa47)* and *egl-8(n488)* PLCβ null mutants compared to wild-type control animals (Fig. 4A-C). Such animals entered infrequent active states but their rhythmic vulval muscle twitching and egg-laying Ca^2+^ transients were grossly intact. In fact, inspection of Ca^2+^ traces suggested an apparent *increase* in vulval muscle Ca^2+^ transient activity in *egl-30(n686)* Gα_q_ and PLCβ mutants (Fig. 4A) including a significant increase in the amplitude and frequency of twitch Ca^2+^ transients in *egl-8(n488)* PLCβ mutant animals (Fig. 4B and 4C). This elevated vulval muscle Ca^2+^ activity was reminiscent of that observed in *egl-1(n986dm)* animals lacking the HSNs (Collins et al., 2016) as these animals still enter and leave infrequent egg-laying active states, possibly driven by the stretch-dependent feedback of egg accumulation in the uterus (Ravi et al., 2018a) which does not appear to act by modulating HSN activity (Ravi et al., 2021). These data further indicate that the egg-laying defects of PLCβ mutants are not caused by a loss of vulval muscle Ca^2+^ activity.

**Figure 4:**
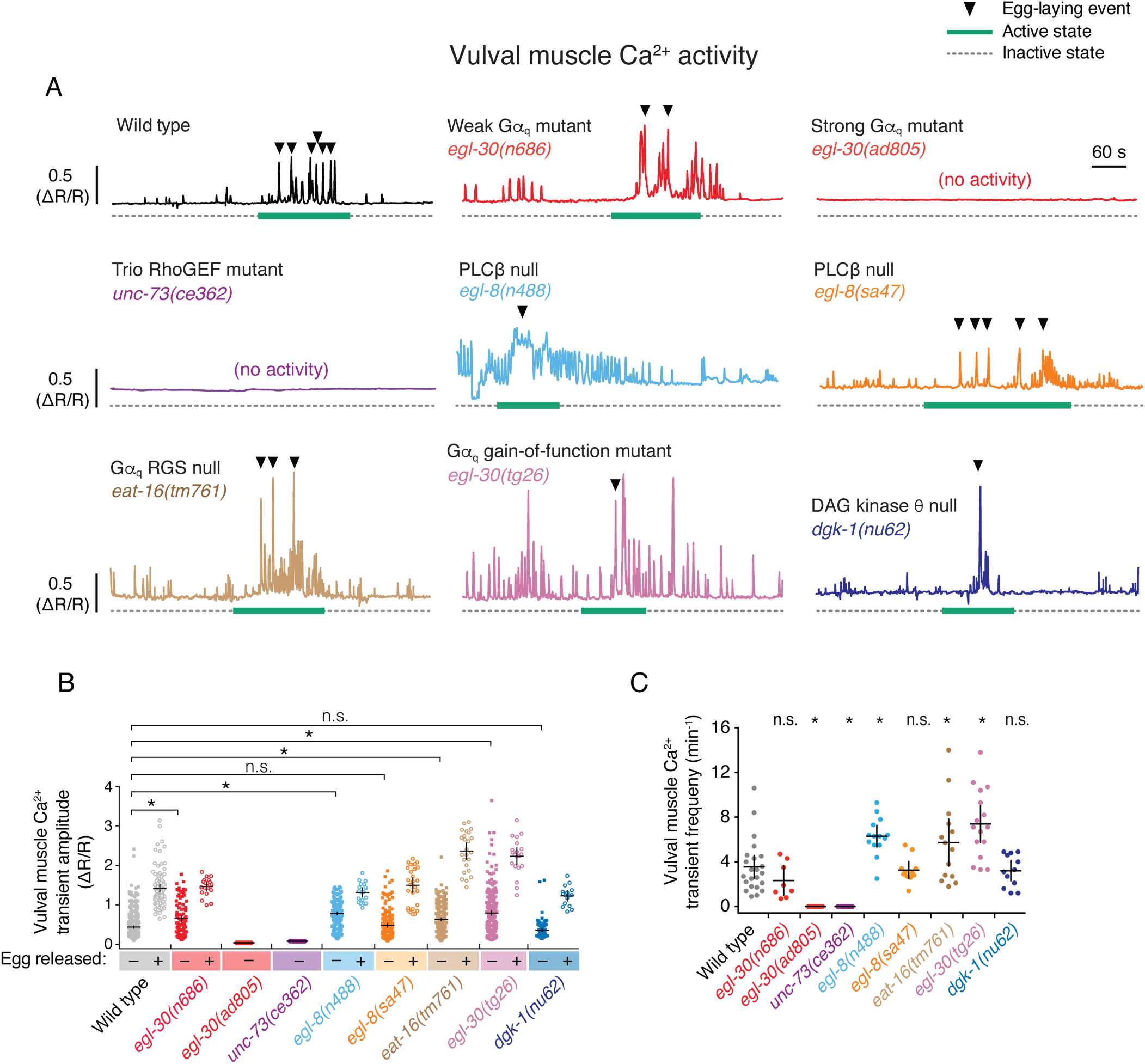
Gα_q_ and Trio signaling promotes vulval muscle activity. **(A)** Representative GCaMP5::mCherry (ΔR/R) ratio traces showing vulval muscle Ca^2+^ activity in freely behaving wild-type (black), *egl-30(n686)* Gα_q_ weak loss-of-function mutant (red), *egl-30(ad805)* Gα_q_ strong loss-of-function mutant (red), *unc-73(ce362)* Trio strong loss-of-function mutant (purple), *egl-8(n488)* PLCβ null mutant (light blue), *egl-8(sa47)* PLCβ null mutant (orange), *eat-16(tm761)* Gα_q_ RGS protein null mutant (grey), *egl-30(tg26)* strong Gα_q_ gain-of-function mutant (magenta), and *dgk-1(nu62)* DAG Kinase null mutant (dark blue) animals during active (green solid bar) and inactive (dotted grey line) egg-laying behavior states. Arrowheads indicate egg-laying events. Vertical and horizontal scale bars show GCaMP5/mCherry fluorescence ratio (ΔR/R) and time, respectively **(B)** Scatterplots of Ca^2+^ transient peak amplitudes for the indicated genotypes during twitch (closed square) and egg-laying transients (open circles). Asterisks indicate p<0.0001, n.s. indicates not significant (p>0.05, Kruskal Wallis test with Dunn’s correction for multiple comparisons). **(C)** Scatterplots of Ca^2+^ transient frequency for indicated genotypes. Line indicates mean eggs laid ±95% confidence intervals; asterisks indicate p≤0.0340; n.s. indicates not significant (p>0.05, One-way ANOVA with Bonferroni’s correction for multiple comparisons; n>10 animals recorded per genotype).

Increased Gα_q_ signaling enhances vulval muscle activity. *egl-30(tg26)* or *eat-16(tm761)* mutant animals with elevated Gα_q_ signaling showed even stronger egg-laying Ca^2+^ transients with average amplitude >2 ΔR/R, a significant difference (Fig. 4A and 4B). Ca^2+^ transients were also significantly more frequent in *egl-30(tg26)* Gα_q_ gain-of-function mutants (∼7 transients per min) and in *eat-16(tm761)* Gα_q_ RGS protein loss-of-function mutants (∼5 transients per min) (Fig. 4C). DAG Kinase-θ (DGK-θ) is thought to antagonize DAG signaling by catalyzing its conversion to phosphatidic acid (Fig. 1A). *dgk-1(nu62)* mutants lacking DGK-θ/DGK-1 have increased neurotransmitter release and egg laying (Jose and Koelle, 2005; Miller et al., 1996; Nurrish et al., 1999), likely through elevation of DAG levels and activation of effectors downstream of Gα_q_. Somewhat surprisingly, *dgk-1(nu62)* mutant animals did not show a significant increase in vulval muscle Ca^2+^ transient amplitude or frequency (Fig. 4A-C). Like PLCβ, DGK-1 is expressed in neurons (Nurrish et al., 1999), suggesting alterations of IP_3_ and/or DAG levels in neurons may affect the frequency of egg-laying active states without altering the overall pattern or strength of vulval muscle Ca^2+^ activity within those active states. Indeed, *goa-1(n1134)* mutants that reduce inhibitory Gα_o_ signaling have hyperactive egg-laying behavior defects that strongly resemble *dgk-1(nu62)* mutants without a significant increase in vulval muscle Ca^2+^ activity (Ravi et al., 2021). Together, these results indicate Gα_q_ and Trio RhoGEF, but not PLCβ, are required for vulval muscle activity that drives twitching and egg-laying Ca^2+^ transients during egg-laying active states.

### DAG mimetics restore muscle activity and egg laying to Gα_q_ signaling mutants

How does serotonin signaling through Gα_q_ promote vulval muscle activity? Previous results have shown that DAG mimetic phorbol esters restore locomotion and animal viability to Gα_q_ null mutants and restore egg laying in PLCβ, Trio RhoGEF double mutants (Williams et al., 2007). However, the effects of phorbol esters on egg laying in single mutants was not clear, raising questions as to whether PMA rescued egg laying downstream of PLCβ, Trio RhoGEF, or both (Fig. 5A). We measured the egg-laying responses of wild-type animals and Gα_q_ signaling mutants to Phorbol 12-myristate 13-acetate (PMA). As shown in Figure 5B, 10 µM PMA treatment strongly stimulated egg laying in wild-type animals and all mutants with reduced Gα_q_ signaling (≥80% animals laying eggs). Like serotonin (Fig. 2B), PMA also rescued egg laying in *egl-1(n986dm)* mutant animals lacking the HSNs (Fig. 5B), but only PMA rescued egg laying in Gα_q_ and effector signaling mutants. These results are consistent with PMA acting downstream of both serotonin release from the HSNs and its subsequent signaling through Gα_q_-coupled receptors. To test whether PMA and DAG act upstream to modulate vulval muscle electrical excitability, we also tested the PMA response in animals carrying mutations in voltage-gated channels that reduce or block egg laying. Loss of L-type Ca^2+^ channel activity in *egl-19(n582)* hypomorphic mutants impairs egg laying downstream of serotonin (Trent et al., 1983; Waggoner et al., 1998), but we find *egl-19(n582)* mutant animals still lay eggs in response to 10 µM PMA (Fig. 5B). The *n582* mutation alters but does not eliminate Ca^2+^ channel function (Gao and Zhen, 2011; Jospin et al., 2002), possibly explaining how PMA could still rescue egg-laying behavior. Indeed, *tpa-1(k530)* mutants originally identified by their resistance to phorbol esters like PMA show synthetic egg-laying defects when combined with *egl-19(n582)* (Waggoner et al., 1998). We next tested gain-of-function K^+^ channel mutants that block egg laying. Animals expressing A383V gain-of-function EGL-23 K2P channels (Ben Soussia et al., 2019; Trent et al., 1983), A331T gain-of-function UNC-103 ERG K+ channels (Collins and Koelle, 2013; Reiner et al., 1999; Reiner et al., 2006), or A478V gain-of-function EGL-2 EAG channels (Weinshenker et al., 1999) showed reduced egg laying in response to PMA (Fig. 5B). PMA-induced egg laying was completely blocked in *egl-23* K2P gain-of-function mutants, and the PMA response was significantly reduced in both *unc-103* ERG and *egl-2* EAG mutant animals compared to wild type (Fig. 5B). Although these results do not rule out that phorbol esters like PMA may be stimulating egg laying in a manner independent of its DAG mimetic effects, our data supports a model where Gα_q_ and Trio RhoGEF signal upstream of DAG to promote vulval muscle excitability and/or contractility.

**Figure 5:**
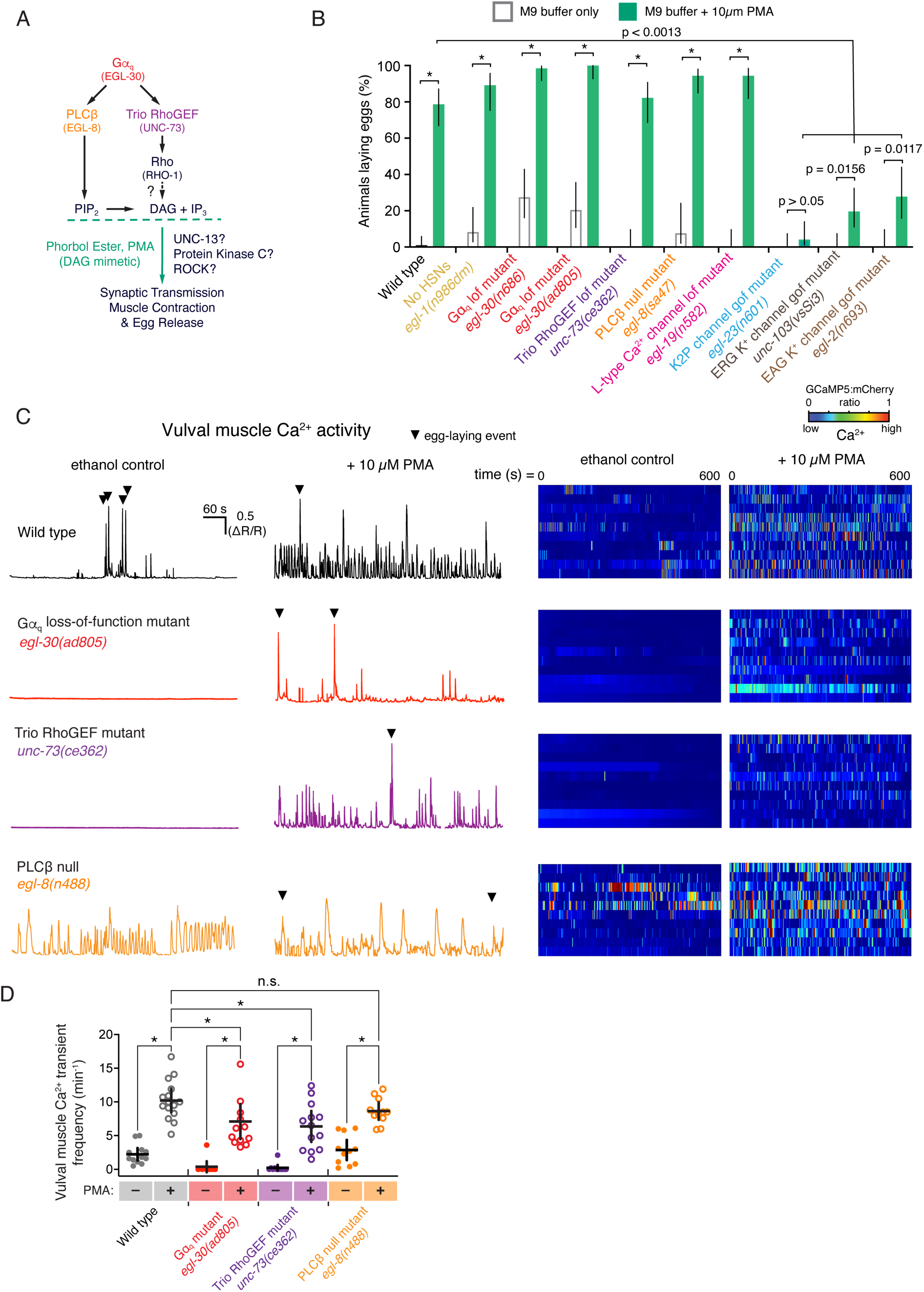
The DAG mimetic PMA rescues egg-laying circuit activity and behavior defects of Gα_q_ signaling mutants. **(A)** Diagrams showing working model of Gα_q_ and DAG signaling pathway during egg-laying behavior. **(B)** Bar plots showing the percentage of animals showing egg laying in M9 buffer (open grey bars) or M9 buffer +10 µM PMA (filled green bars). Error bars indicate 95% confidence intervals for the proportion; asterisks indicate p<0.0013 (Fisher’s exact test with Bonferroni’s correction for multiple comparisons; n≥30 animals per genotype and condition). **(C)** Left, representative GCaMP5::mCherry (ΔR/R) ratio traces showing vulval muscle Ca^2+^ activity in wild-type or the indicated Gα_q_ signaling mutant animals in the absence or presence of 10 µM PMA. Arrowheads indicate egg-laying events. Vertical and horizontal scale bars show GCaMP5/mCherry fluorescence ratio (ΔR/R) and time, respectively. Right, heat map showing intensity modulated color spectrum of GCaMP5::mCherry (ΔR/R) ratio of vulval muscle Ca^2+^ activity ranging from blue (low Ca^2+^) to red (high Ca^2+^). Rows indicate ratio changes in each of 10 animals. **(D)** Scatterplots of Ca^2+^ transient frequency in the absence (-) and presence (+) of 10 µM PMA for the indicated genotypes. Lines indicate mean eggs laid ±95% confidence intervals; asterisk indicates p≤0.0275; n.s.= not significant (p>0.05, One-way ANOVA with Bonferroni’s correction for multiple comparisons; n≥10 animals per genotype and condition).

We next imaged how PMA affected vulval muscle Ca^2+^ activity. We performed 10-minute GCaMP5 Ca^2+^ recordings of wild-type or Gα_q_ signaling mutants after two hours of exposure to 10 µM PMA (Movies 1-3). Quantitation of Ca^2+^ transients showed that PMA significantly increased vulval muscle Ca^2+^ activity in wild-type animals to 10±1.6 transients per minute from an average 2±0.9 transients per min in vehicle-treated, wild-type animals (Fig. 5C-D and Movie 1). PMA also restored both rhythmic twitch and egg-laying Ca^2+^ transients to strong Gα_q_ and Trio RhoGEF signaling mutants, with Ca^2+^ transients frequencies increasing from essentially 0±0.8 transients per minute in vehicle-treated controls to 7±2 transients per minute after PMA treatment, almost but not quite to the level of PMA-treated wild-type control animals (Fig. 5C-D and Movies 2-3). Together, these studies show that DAG-mimetic phorbol esters restore muscle excitability and contractility defects of Gα_q_ and Trio RhoGEF mutants, suggesting DAG production could be a major and necessary consequence of both Gα_q_ and Trio RhoGEF signaling.

### Phorbol esters promote egg laying independent of UNC-13 or single Protein Kinase C isoforms

How do phorbol esters like PMA rescue vulval muscle Ca^2+^ activity and egg laying? Previous results have shown that DAG and PMA bind to C1 domain-containing proteins such as mUNC-13/UNC-13 and Protein Kinase C (PKC) to regulate their activity (Betz et al., 1998; Huang, 1989; Konig et al., 1985; Newton, 2001; Silinsky and Searl, 2003).To test if DAG regulates egg-laying behavior through activation of UNC-13 or PKC, we tested whether mutants lacking these proteins still have a robust serotonin and/or PMA egg-laying response (Fig. 6). Mutants that eliminate axonal and synaptic UNC-13 show reduced egg laying, accumulating an average of 22 eggs compared to 15 seen in wild-type animals (Fig. 6A), like PLCβ mutants but significantly fewer than the >30 eggs that accumulate in Gα_q_ and Trio RhoGEF mutants (Fig. 1C-I). *unc-13* mutants also resemble PLCβ mutants in their egg-laying response to serotonin and PMA. Egg laying in *unc-13* mutants was stimulated by PMA but was resistant to serotonin (Fig. 6B). While the serotonin resistance we observed for *unc-13* mutants after one hour differs from that seen by Bastiani et al., 2003 at 90 minutes, we saw similar differences for the PLCβ mutants (Table 3). These results show that serotonin promotes egg laying through a PLCβ and UNC-13-dependent pathway that may be distinct from the PMA-stimulated pathway.

**Figure 6:**
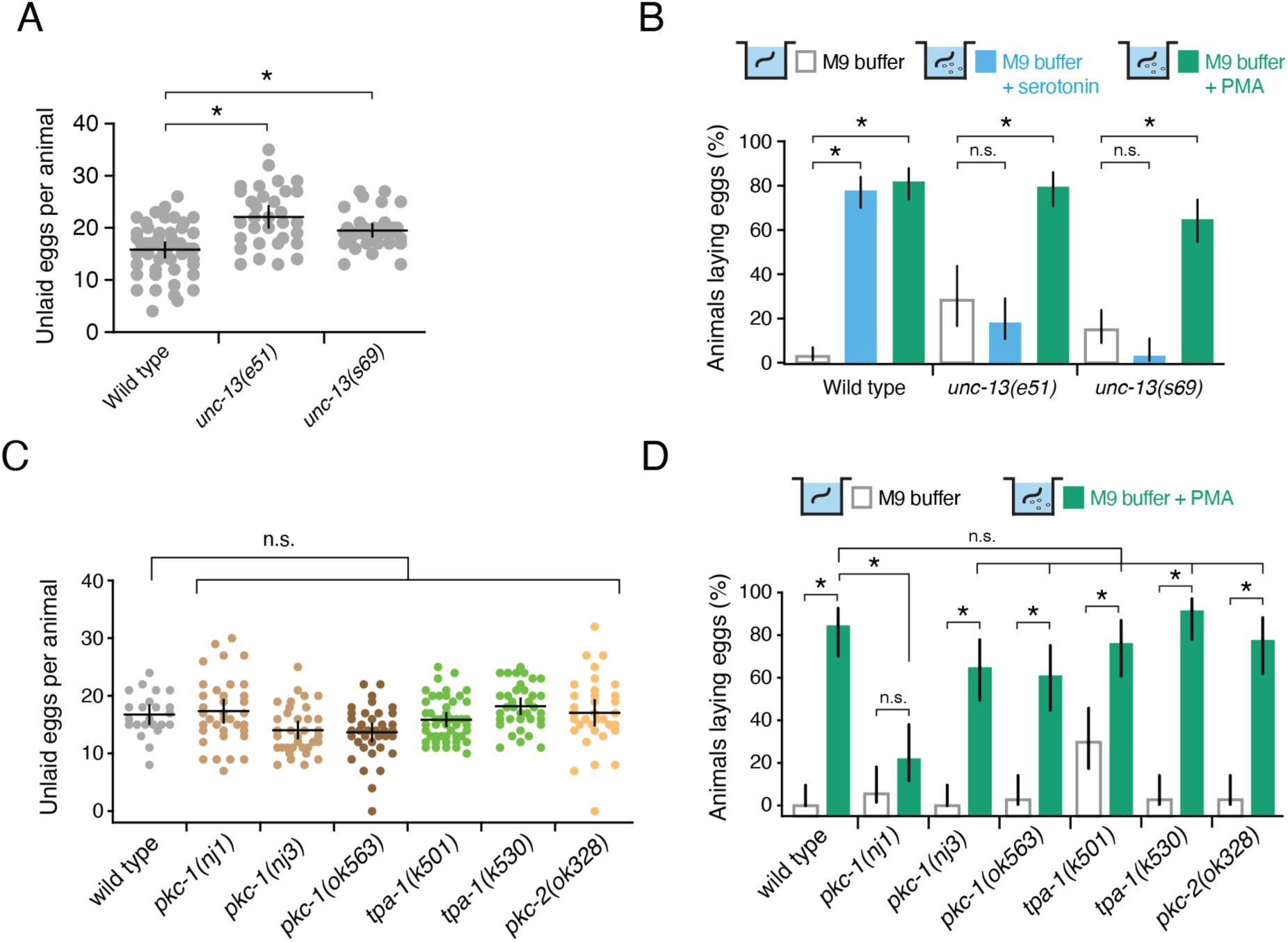
DAG promotes egg laying independent of UNC-13 or Protein Kinase C. **(A)** Scatterplot of egg accumulation in wild-type, *unc-13(e51)* loss-of-function mutant, and *unc-13(s69)* null mutant animals. Lines indicate mean eggs laid ±95% confidence intervals. Asterisk indicates p≤0.0021 (One-way ANOVA with Bonferroni’s correction for multiple comparisons; n≥36 per genotype). (**B)** Bar plots showing the percentage of wild-type, *unc-13(e51),* or *unc-13(s69)* mutant animals laying eggs in M9 buffer (grey open boxes), 18.5 mM serotonin (filled blue boxes), or 10 µM PMA (filled green boxes). Asterisks indicate p<0.0006; n.s. = not significant (p>0.05, Fisher’s exact test with Bonferroni’s correction for multiple comparisons; n≥36 animals per genotype and condition). **(C)** Scatterplot of egg accumulation in wild-type (n=24) and the indicated Protein Kinase C mutant animals (n≥35 per genotype). Line indicates mean eggs accumulated ±95% confidence intervals. n.s.= not significant (p>0.05, One-way ANOVA with Bonferroni’s correction for multiple comparisons. **(D)** Bar plots showing the percentage of wild-type and Protein Kinase C mutant animals showing egg laying in M9 buffer (open bars) or 10 µM PMA (filled green bars). Bar indicates mean eggs ±95% confidence intervals for the proportion. Asterisks indicate p<0.0007; n.s.= not significant (p>0.05, Fisher’s exact test with Bonferroni correction for multiple comparisons; n≥35 animals per genotype and condition).

To determine whether Gα_q_ and Trio RhoGEF signaling through the PMA-responsive pathway is mediated by Protein Kinase C, we analyzed egg accumulation in animals bearing predicted null mutations in different PKC isoforms (Edwards et al., 2012; Hyde et al., 2011; Okochi et al., 2005; Tabuse, 2002; Tabuse et al., 1989). The *C. eleg*ans genome encodes four PKCs isoforms PKC-1, PKC-2, PKC-3, and TPA-1. PKC-1 has previously been shown to promote neuropeptide transmission (Sieburth et al., 2007). While neuropeptides signal to promote egg laying (Avery et al., 1993; Brewer et al., 2019; Jacob and Kaplan, 2003; Kass et al., 2001), PKC-1 (nPKC-ε) null mutants showed a grossly normal egg accumulation of 14∼17 eggs (Fig. 6C). Animals bearing predicted null mutants of novel and conventional PKCs such as nPKCδ/θ (TPA-1) and cPKCα/β (PKC-2) orthologs also show no significant differences in egg accumulation (Fig. 6C), suggesting that, unlike loss of Gα_q_ or Trio RhoGEF signaling, disruption of individual PKC signaling pathways do not strongly affect egg-laying behavior. We next tested whether PKC mediates the egg-laying response to PMA. All the PKC single mutant animals laid eggs in response to PMA except for *pkc-1(nj1)* (Fig. 6C). The *nj1* allele is predicted to be a missense mutation that may lead to the expression of a mutant PKC protein with altered function (Okochi et al., 2005). Indeed, previous experiments using this mutant have shown that *pkc-1(nj1)* mutant animals have stronger behavior defects (Okochi et al., 2005; Ventimiglia and Bargmann, 2017), suggesting the mutant protein may be expressed and interfere cell signaling, possibly by interfering with the function of other co-expressed PKC isoforms like TPA-1. Together, these results support a model where Gα_q_ and Trio RhoGEF signaling in the vulval muscles drives elevation of DAG which activates targets like PKCs and/or other effectors to promote cell electrical excitability for egg laying.

## Discussion

In this study we explored the cellular and molecular specificity of Gα_q_ effector signaling as it regulates egg-laying circuit activity and behavior using molecular genetics, optogenetics, pharmacology, and Ca^2+^ imaging techniques. We found that Gα_q_ effectors PLCβ and Trio RhoGEF differentially act in neurons and muscles to promote synaptic transmission and egg-laying behavior, supporting a working model where Gα_q_ signals through Trio RhoGEF in both neurons and muscles while PLCβ functions outside of HSN to promote egg laying (Fig. 7). Although Gα_q_, PLCβ, and Trio RhoGEF mutants all fail to lay eggs in response to serotonin, optogenetic stimulation of HSNs fully rescued egg laying in PLCβ but not Gα_q_ or Trio RhoGEF mutants. Recent work has shown that the HSNs release NLP-3 neuropeptides which can promote egg laying even in *tph-1* mutants lacking serotonin (Brewer et al., 2019), possibly explaining why optogenetic activation of HSNs rescues egg laying to PLCβ mutants even when exogenous serotonin cannot. The HSNs are also predicted to release ACh (Pereira et al., 2015), and nAChR receptors are expressed on the vulval muscles and can stimulate egg laying (Kim et al., 2001; Waggoner et al., 2000). Because serotonin drives egg laying in animals lacking HSNs but not in animals lacking PLCβ, and that expression of PLCβ is sufficient to rescue normal egg laying, we propose working model (Fig. 7) where Gα_q_ and PLCβ act in neurons other than HSNs to promote release of neurotransmitters like ACh onto the vulval muscles to stimulate egg laying. The cholinergic VA, VB, and VC neurons also innervate the vulval muscles alongside the HSNs and express PLCβ (Cook et al., 2019; Taylor et al., 2021; White et al., 1986). Consistent with this model, we have recently shown that blocking VC synaptic transmission reduces egg laying in response to serotonin (Kopchock et al., 2021). Optogenetic stimulation of the VCs (Kopchock et al., 2021) or VA/VB neurons (Kopchock, 2021) stimulates vulval muscle Ca^2+^ activity, but the resulting Ca^2+^ activity is insufficient to drive the strong egg-laying contractions. Because we find that mutations that increase Gα_q_ signaling have stronger vulval muscle Ca^2+^ transients, we propose that serotonin and NLP-3 released from HSN potentiates ACh release from other motor neurons and promotes the electrical excitability and/or contractility of the vulval muscles, converting rhythmic twitch Ca^2+^ transients into stronger egg-laying transients. Previous studies have shown that Trio RhoGEF acts in neurons to regulate locomotion behavior (Hu et al., 2011; Steven et al., 2005; Williams et al., 2007). Our studies show that transgenic Trio RhoGEF expression in either neurons or muscles alone is insufficient to restore wild-type level of egg-laying behavior, but expression in both is sufficient. These results mirror previously published results for Gα_q_ (Bastiani et al., 2003), further supporting a model where PLCβ and Trio RhoGEF functions during locomotion and egg-laying behaviors are distinct.

**Figure 7:**
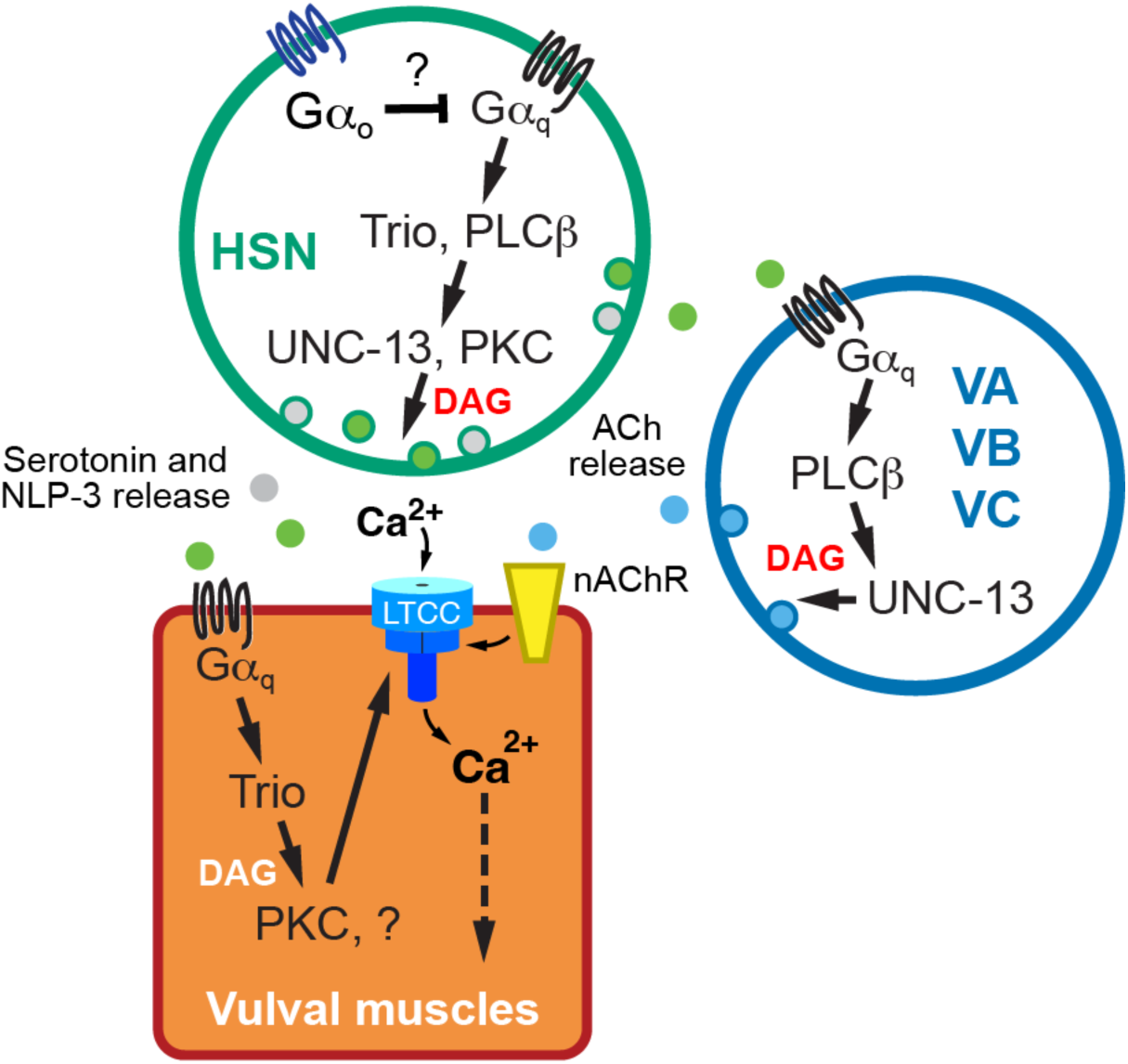
Working model of Gα_q_ signaling in the egg-laying circuit. See text for details.

How does Gα_q_, PLCβ, and Trio-RhoGEF signaling promote egg-laying behavior? Earlier studies have suggested that Rho ortholog RHO-1 in *C. elegans* regulates synaptic activity in a mechanism that involves the G12 family protein GPA-12 (Hiley et al., 2006; Lutz et al., 2005). Activated RHO-1 also directly binds to and inhibits the DGK-1 diacylglycerol kinase expressed in neurons that signals to reduce DAG available to bind effectors (Hiley et al., 2006; McMullan et al., 2006). Our data are consistent with the previous results reporting that Gα_q_ signaling regulates postsynaptic vulval muscle activity mainly through Gα_q_-Trio pathway as Gα_q_ and Trio mutants show a similarly strong reduction in vulval muscle Ca^2+^ activity. Because muscle activity defects in Gα_q_ and Trio mutants can be restored by the DAG-mimetic PMA, we suggest that insufficient levels of DAG are responsible for the circuit activity and behavior defects of Gα_q_ and Trio RhoGEF mutants. In the absence of PLCβ, how would parallel Gα_q_ signaling through Trio RhoGEF and RHO-1 generate DAG? Besides PLCβ (EGL-8)*, C. elegans* expresses four other PLC orthologs: PLCε (PLC-1), PLC-2, PLCγ (PLC-3), and PLCδ (PLC-4) (Vázquez-Manrique et al., 2008). *In vitro* studies with cultured mammalian cells show that small G proteins like Rho can bind to and activate PLCε (Seifert et al., 2008; Wing et al., 2003). Genetic and molecular expression evidence in *C. elegans* suggests a model where PLCε is activated downstream of Gα_q_ and Rho to promote cell activity (Kunitomo et al., 2013; Taylor et al., 2021; Yu et al., 2013), but whether Trio activation of Rho ultimately acts through these PLCs to produce DAG is not clear. One approach to test if these other PLCs mediate Rho signaling would be to perform genetic epistasis experiments. However, loss of Rho-1 causes lethality (Jantsch-Plunger et al., 2000; McMullan and Nurrish, 2011) and loss of PLCε cause sterility defects (Yin et al., 2004), limiting our ability to measure differences in egg laying. Alternatively, Gα_q_ signaling through the Rho-1 branch may be independent of PLCs and DAG production where exogenous PMA is instead activating factors downstream of a parallel PLCβ pathway. While our rescue data are consistent with PLCβ acting in neurons and Trio RhoGEF acting in muscles, we cannot rule out a PLCβ function for DAG or IP_3_ production in the vulval muscles. Imaging or biochemical approaches documenting Gα_q_-dependent changes in PIP_2_ (Stauffer et al., 1998) and/or DAG (Ohno et al., 2017; Tewson et al., 2012) *in vivo*, along with cell-specific rescue and knockout experiments (LeBoeuf et al., 2020), should resolve whether the Rho-1 branch acts through PLCs and/or inhibits DAG lipases to promote DAG levels.

Do phorbol esters like PMA stimulate *C. elegans* egg laying by acting as DAG mimetics? Previous studies have shown that phorbol esters promote both synaptic vesicle and dense core vesicle release from neurons and neurosecretory cells (Silinsky and Searl, 2003). DAG and phorbol esters activate many effectors including mUNC-13 in the brain and Protein Kinase C (PKC) in nearly all cells (Betz et al., 1998; Huang, 1989). In *C. elegans*, exogenous treatment with phorbol esters causes growth inhibition, uncoordinated movement, and lethality which can be suppressed by loss-of-function mutations in a single gene, *tpa-1*, which encodes nPKCδ/θ (Tabuse and Miwa, 1983; Tabuse et al., 1989). Acute PMA treatment promotes hypersensitivity to the paralytic effects of aldicarb (Sieburth et al., 2007). Double mutants of both PKC-1 and UNC-13 (H17K) show increased resistance to phorbol esters compared to either mutant alone suggesting that PMA acts in part through these effectors to regulate ACh release (Sieburth et al., 2007; Silinsky and Searl, 2003). Our data show that animals lacking UNC-13 still lay eggs in response to PMA. Mutant animals with defects in single PKC isoform encoding genes were similarly responsive to PMA, with the notable exception of *pkc-1(nj1)* mutant animals. *pkc-1(nj1)* results in a missense mutation and shows a significantly reduced PMA response compared to two other putative null mutants. Such allele-specific differences among *pkc-1* alleles have been observed previously in experiments studying PKC function in nose touch response, octanol and high osmolarity avoidance (Hyde et al., 2011), regulation of AWC^ON^ glutamate release (Ventimiglia and Bargmann, 2017), and in regulation of PKC by DAG or Ca^2+^ (Okochi et al., 2005). The *nj1* allele may impart a dominant-negative effect, affecting the recruitment or function of other PKC isoforms. For example, TPA-1 has been shown to function redundantly with PKC-1 (also known as TTX-4), a nPKC-ε ortholog (Okochi et al., 2005). Studies have shown that activated Gα_q_ and accumulation of DAG recruit TPA-1 to compensate for the loss of PKC-1 (Hiroki and Iino, 2022). Egg-laying defects of *egl-19(n582)* L-type Ca^2+^ channel mutants are enhanced when combined with *tpa-1(k530)* PKC null mutants (Waggoner et al., 1998), suggesting TPA-1 may mediate some of the DAG and/or PMA response for egg laying. Future work testing compound mutants disrupting UNC-13 and different PKC isoforms should reveal whether PMA acts as a DAG mimetic in neurons and muscle cells to promote egg laying.

Besides the compensatory effect of various PKCs and UNC-13, this study does not rule out other effectors as potential targets of DAG and/or PMA. *In vitro* studies show ROCK (Rho-associated coiled-coil kinase) activation in PMA-induced apoptosis and macrophage differentiation (Chang et al., 2006; Yang et al., 2017). ROCK has a predicted C1 domain that mediates protein interaction with DAG and might bind and be similarly activated by PMA (Xiao et al., 2009). In *C. elegans,* RHO-1 signals through LET-502/ROCK to phosphorylate non-muscle myosin light chain (Shimizu et al., 2018). Gα_q_ also promotes neurotransmitter release via additional kinase targets including SEK-1 Mitogen-Activated Protein Kinase Kinase in the p38 MAPK pathway and KSR-1 in the ERK MAPK pathway (Coleman et al., 2018; Hoyt et al., 2017). KSR-1 is particularly interesting in that its N-terminus shares sequence similarly with C1 domains that might mediate regulation by DAG. Loss of KSR-1 and other ERK MAPK components also suppress the loopy locomotion defects caused by gain-of-function mutations in Rho-1 (Coleman et al., 2018). Taken together, our work is consistent with a model where additional DAG-sensitive effectors act downstream of Gα_q_, Trio, and Rho to promote muscle excitability and /or contractility for egg laying.

Phorbol esters and locally produced DAG may promote egg laying via activation of distinct effectors. Apart from activation of C1 domain containing effectors, emerging evidence indicates Phospholipase C-dependent production of DAG directly modulates the gating of ion channels for membrane excitability. For example, DAG activates several ion channels including canonical transient receptor potential (TRP) cation channels (TRPC) (Hofmann et al., 1999) while also inhibiting other ion channels including two-pore domain TASK potassium channels via an unknown mechanism (Wilke et al., 2014). PIP_2_ has also been shown to modulate some ion channels like KCNQ directly (Suh et al., 2006), although DAG and PMA modulate *C. elegans* KCNQ channels likely via the intermediate activation of protein kinases including PKC (Wei et al., 2005). Thus, Gα_q_ modulation of PIP_2_ and DAG levels could directly or indirectly affect several postsynaptic ion channels to shape electrical excitability. DAG is also precursor in the production of several signaling lipids, including the endocannabinoid 2-arachidonoylglycerol (2-AG) which has been shown to signal from dendrites in a retrograde manner through neuronal Gα_o_-coupled endocannabinoid receptors to inhibit neurotransmitter release (Hashimotodani et al., 2013; Hashimotodani et al., 2005; Soltesz et al., 2015; Tanimura et al., 2010; Wettschureck et al., 2006). In *C. elegans*, 2-AG activates the NPR-19 endocannabinoid receptor ortholog that couples to Gα_o_ to modulate serotonin transmission, pharyngeal, feeding, and locomotory behaviors (Oakes et al., 2019; Oakes et al., 2017; Pastuhov et al., 2016). We have recently shown that feedback of egg accumulation alters vulval muscle Ca^2+^ activity, which subsequently signals to regulate bursts of Ca^2+^ transients in the HSNs that accompany the onset of the egg-laying active state (Ravi et al., 2018a; Ravi et al., 2021). These results support a model where stretch-dependent feedback of egg accumulation stimulates postsynaptic vulval muscle Ca^2+^ signaling. This Ca^2+^ could then activate PLCs to generate DAG and 2-AG which signal to modulate HSN activity, serotonin release, and egg laying. The genetic and experimental accessibility of the *C. elegans* egg-laying circuit should allow us to determine if conserved G proteins like Gα_q_ act generally to drive neural circuit activity via changes in DAG, subsequent activation of effectors, and retrograde messengers like 2-AG.

## Acknowledgements

This work was funded by grants from the NIH (R01-NS086932) and NSF (IOS-1844657) to KMC. PD is supported by AHA predoctoral fellowship Award (20PRE35210233). We thank Drs. Kenneth Miller, Joshua Kaplan, and Thomas Boulin for sharing plasmids and strains. Some of the strains used in this study were provided by the *C. elegans* Genetics Center, which is funded by NIH Office of Research Infrastructure Programs (P40 OD010440). We thank Drs. Julia Dallman, Laura Bianchi, and Athula Wikramanayake along with members of the Collins lab for helpful discussions and feedback on the manuscript.

---

**Movie 1: GCaMP5:mCherry fluorescence ratio in the vulval muscles of wild-type animals in the absence or presence of 10 µM DAG analogue, PMA.** Blue indicates low Ca^2+,^ and red indicates elevated Ca^2+^.

**Movie 2: GCaMP5:mCherry fluorescence ratio in the vulval muscles of strong loss of function Gα_q_ mutant, *egl-30(ad805),* animals in the absence or presence of 10 µM DAG analogue, PMA.** Blue indicates low Ca^2+,^ and red indicates elevated Ca^2+^.

**Movie 3: GCaMP5:mCherry fluorescence ratio in the vulval muscles of strong loss of function Trio mutant, *unc-73(ce362)*, animals in the absence or presence of PMA.** Blue indicates low Ca^2+,^ and red indicates elevated Ca^2^.

